# Molecular layer interneurons in the cerebellum encode for valence in associative learning

**DOI:** 10.1101/2019.12.14.876201

**Authors:** Ming Ma, Gregory L. Futia, Fabio M. Simoes De Souza, Baris N. Ozbay, Isabel Llano, Emily A. Gibson, Diego Restrepo

## Abstract

We used two-photon microscopy to study the role of ensembles of cerebellar molecular layer interneurons (MLIs) in a go-no go task where mice obtain a sugar water reward if they lick a spout in the presence of the rewarded odorant and avoid a time out when they refrain from licking for the unrewarded odorant. In naïve animals the MLI responses did not differ between the odorants. With learning, the rewarded odorant elicited a large increase in MLI calcium responses, and the identity of the odorant could be decoded from the differential response. Importantly, MLIs switched odorant responses when the valence of the stimuli was reversed. Finally, mice took a longer time to refrain from licking in the presence of the unrewarded odorant and had difficulty becoming proficient when MLIs were inhibited by chemogenetic intervention. Our findings support a role for MLIs in learning valence in the cerebellum.

---

The cerebellum plays a pivotal role in coordinating movements through sensorimotor integration. It receives massive input through mossy fiber (MF) synapses onto granule cells (GCs) to form a circuit efficient in complex pattern separation^1^ (Supplementary Fig. 1). In addition, the cerebellum contributes to cognition and emotion and is associated with non-motor conditions such as autism spectrum disorders^2-5^. The Purkinje cells (PCs), the sole projection neurons of the cerebellar cortex, receive excitatory afferent input from the parallel fibers (PFs) of GCs and dense feedforward inhibitory inputs from the molecular layer interneurons (MLIs). Importantly, subtle changes in this PC excitatory-inhibitory balance generate robust, bidirectional changes in the output of PCs^6^.

Plasticity in cerebellar circuit activity plays an important role in generation of adequate output. Indeed, long term depression (LTD) mediated by dendritic increases in Ca^2+^ in PCs elicited by motor error signals conveyed by climbing fibers (CFs) is a classical model of plasticity^7-10^. However, recent studies indicate that CFs also signal reward prediction^11,12^ or decision-making errors^13^, and the cerebellum modulates association pathways in the ventral tegmental area (VTA) contributing to reward-based learning and social behavior^14^. Furthermore, although LTD at the PF-PC synapse is often considered as the substrate for cerebellar dependent learning^7-10^, such learning can occur in the absence of LTD and may therefore involve other forms of plasticity^15^. A potential substrate for plasticity is the PF-MLI synapse^16,17^ where LTP can be induced in slices by pairing MLI depolarization with PF stimulation^18^ and *in vivo* by conjunctive stimulation of PFs and CFs^19^, believed to underlie changes in the size of cutaneous receptive fields^20,21^. Additionally, high frequency stimulation of PFs alters subunit composition of AMPA receptors, rendering them calcium-impermeable^18^, a long-lasting change linked to behavioral modifications^22^. Interestingly, MLIs have been proposed to participate in cerebellar plasticity^21,23,24^, and Rowan and co-workers found graded control of PC plasticity by MLI inhibition^25^ suggesting that MLI inhibition is a gate for learning stimulus valence which conveys information as to whether the stimulus is rewarded. However, whether there is a causal participation of MLIs in reward-associated learning is unknown.

Here we explored whether MLI activity plays a role in reward-associated learning in a go-no go task where the thirsty animal learns to lick to obtain a water reward^26,27^. We applied two-photon microscopy^28^ to record Ca^2+^ changes in ensembles of MLIs and utilized chemogenetics to explore the functional role of MLI activity in learning.

## Results

In order to explore the role of MLIs in associative learning we employed *in vivo* two-photon microscopy to record neural activity reported by changes in fluorescence emitted by the Ca^2+^ indicators GCaMP6/7. MLIs were imaged within the superficial 50 μm of the ML in head-fixed mice through a 2×2 mm glass window implanted above the cerebellar vermis (lobule VI, Fig. 1, Supplementary Fig. 2), where GCs acquire a predictive feedback signal or expectation reward^29,30^. We used the go-no go task where thirsty mice initiate the trial by licking on the spout to elicit odorant delivery 1-1.5 sec after the first lick. Mice received a water reward when they licked at least once in two lick segments during rewarded odorant delivery (1% isoamyl acetate, Iso, termed S+) (Fig. 1a, Hit trial, mouse movement shown in Supplementary Movie 1, quantified in Supplementary Fig. 3). Mice did not receive the reward if they failed to lick in one of the two lick segments (Miss trial). When the unrewarded odorant was presented (mineral oil, MO, S-trials) the animals did not receive a reward when they refrained from licking (Correct Rejection, CR, Fig. 1a), and they received a timeout of ten seconds if they licked in both segments (False Alarm, FA). A proficient mouse (percent correct>=80%) licked at least once in each lick segment in the S+ trials and stopped licking during the segments for the S-trials, see lick trace at the bottom of Fig. 1d.

**Figure 1.**
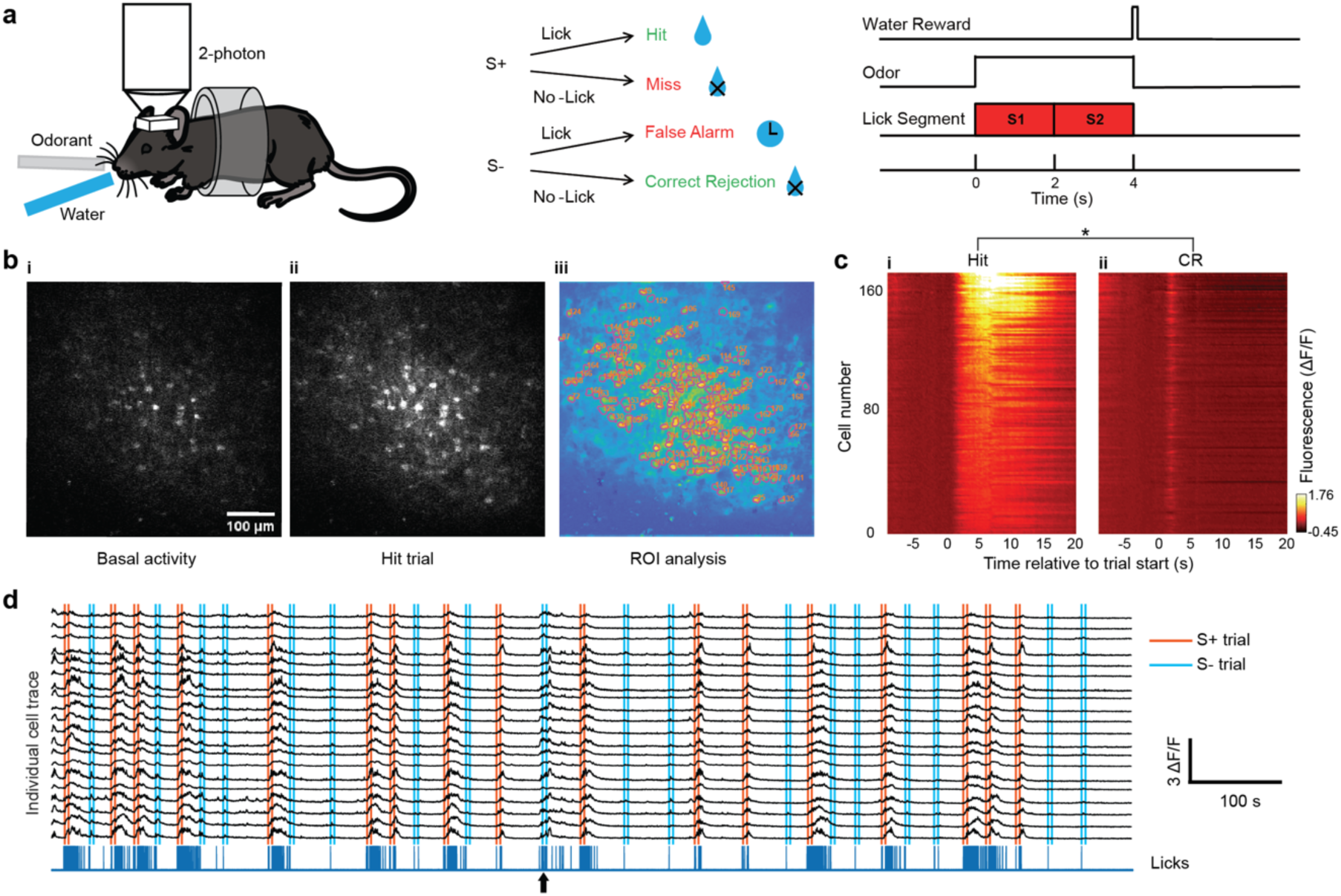
Two-photon Ca^2+^ imaging of MLIs in head-fixed mice undergoing the go-no go associative learning task. **a**. Go-no go task. Left: Two-photon imaging of a head-fixed mouse responding to odorants by licking on a water spout in response to the rewarded odorant in the go-no go task. Center: Scoring of decision making. Right: time course of the trial. For water reward in Hit trials the animal must lick at least once in each of the two 2 second lick segments. **b**. The panels in **i** and **ii** show two photon microscopy images of GCaMP6f fluorescence recorded from MLIs in a mouse proficient in the go-no go task (basal activity: before trial start, Hit trial: during reinforced odorant application). The color image in **iii** shows the regions of interest identified using CaImAn software^31^. **c**. Pseudocolor plots displaying the average per trial ΔF/F time course for Hit (**i**) and CR (**ii**) odorants for all ROIs in this example. GLM analysis involving time periods pre-odorant (1 sec before odorant onset), odorant (last second during odorant application) and reinforcement (1.5 second after reinforcement) and different events (Hits, Miss, CR and FA) yielded significant differences between reinforcement and pre-odorant (p<0.001), between odorant and pre-odorant (p<0.001), and between all interactions between these two period pairs and all events (p<0.01, 2040 observations, 2028 degrees of freedom, n=170 ROIs, 1 mouse, GLM F-statistic 234, p<0.001). *Post-hoc ranksum p<pFDR=0.048. **d**. Neural ensemble activity. Black traces are the GCaMP6f fluorescence (ΔF/F) time courses for a subset of the ROIs identified in the FOV in **b**. This mouse was proficient (>=80% correct trials). Vertical lines: orange start and end of S+ odorant application, light blue is S-. The blue trace at the bottom shows the licks. All trials were Hits or CRs with the exception of the trial identified with the arrow that was a FA. Data shown in panels b-d are from one session (one mouse).

An example of an increase in MLI GCaMP6f fluorescence intensity during odorant application in a Hit trial is shown for a proficient mouse in Figs. 1bi,ii (Supplementary Movie 2). We extracted regions of interest (ROIs) of the components (Fig. 1biii) and temporal traces of the normalized change in fluorescence (ΔF/F, Figs. 1c and d) using constrained nonnegative matrix factorization analysis^31^. The average diameter of the ROIs in this field of view (FOV) was 10.5±4 μm (mean±SD, n=170, Supplementary Fig. 4a), which falls within the range of diameters reported for MLIs^32^. When the animal was proficient we observed that MLI ΔF/Fs increased when the rewarded odorant was presented (Fig. 1ci and d, vertical orange lines in 1d indicate odorant on and off times for S+). In contrast, the unrewarded odorant elicited smaller transient increases in ΔF/F (Fig. 1cii, Fig. 1d vertical light blue lines and Supplementary Figs. 4b,c,d), with some exceptions where the increases for ΔF/F for S-were larger (arrow in Fig. 1d). Finally, the rewarded odorant elicited increases in ΔF/F in 89% of the ROIs among a total of 191 (Fig. 1c and Supplementary Figs. 4c and d).

### Cerebellar MLIs develop divergent responses as mice learn to differentiate odorants

The behavioral performance in a learning session where the animal reached proficient level after 70 trials is shown in Fig. 2a. When the animal was naïve (<=65% correct) the time courses for ΔF/F overlapped between S+ and S-(Fig. 2bi), and when the animal became proficient the time courses for ΔF/F diverged (Fig. 2biii). For the proficient animal in S+ trials ΔF/F started increasing at trial initiation (1-1.5 sec before odorant addition) and reached a peak when the animal was rewarded (Fig. 2biii, orange trace), while for S-trials, the response returned to basal levels shortly after the odorant was applied (Fig. 2biii, light blue trace, Fig. 2b shows the time courses for lick rates). Whereas ΔF/F does not differ between S+ and S-before odorant addition (Fig. 2c), it diverges after odorant addition as the animal learns (Fig. 2d) and ΔF/F per ROI increased when the animal became proficient (Fig. 2e). GLM analysis yields a statistically significant difference between S+ and S- and naïve and proficient, p<0.001, 5550 observations, 5538 d.f., 1 mouse, GLM F-statistic=267, p<0.001. Finally, the mean ΔF/F (±CI, 4 mice) for the last second of odorant application increases as a function of percent correct performance for the S+ condition whereas it remains stable for the S-condition (Fig. 2g). A GLM analysis finds a statistically significant difference between S+ and S-(p<0.01, 47 observations, 44 d.f., n=4 sessions, 4 mice, GLM F-statistic=60, p<0.001) and for the interaction between performance and the odorant (S+ vs. S-) (p<0.001, 47 observations, 44 d.f., n=4 sessions, 4 mice (Supplementary Fig. 6 shows learning curves for these mice). Therefore, as the animal learns, the magnitude and temporal dynamics of the change in Ca^2+^ in MLIs diverges between S+ and S-odorants, suggesting a possible role for MLIs in associative learning. Interestingly, the majority of the ROIs responded similarly (Fig. 1c, Supplementary Fig. 4) and a dimensionality analysis indicated that the dimensionality ranged from 2 to 6 (see Supplementary Note 1 and Supplementary Fig. 7) indicating that the responses are highly redundant.

**Figure 2.**
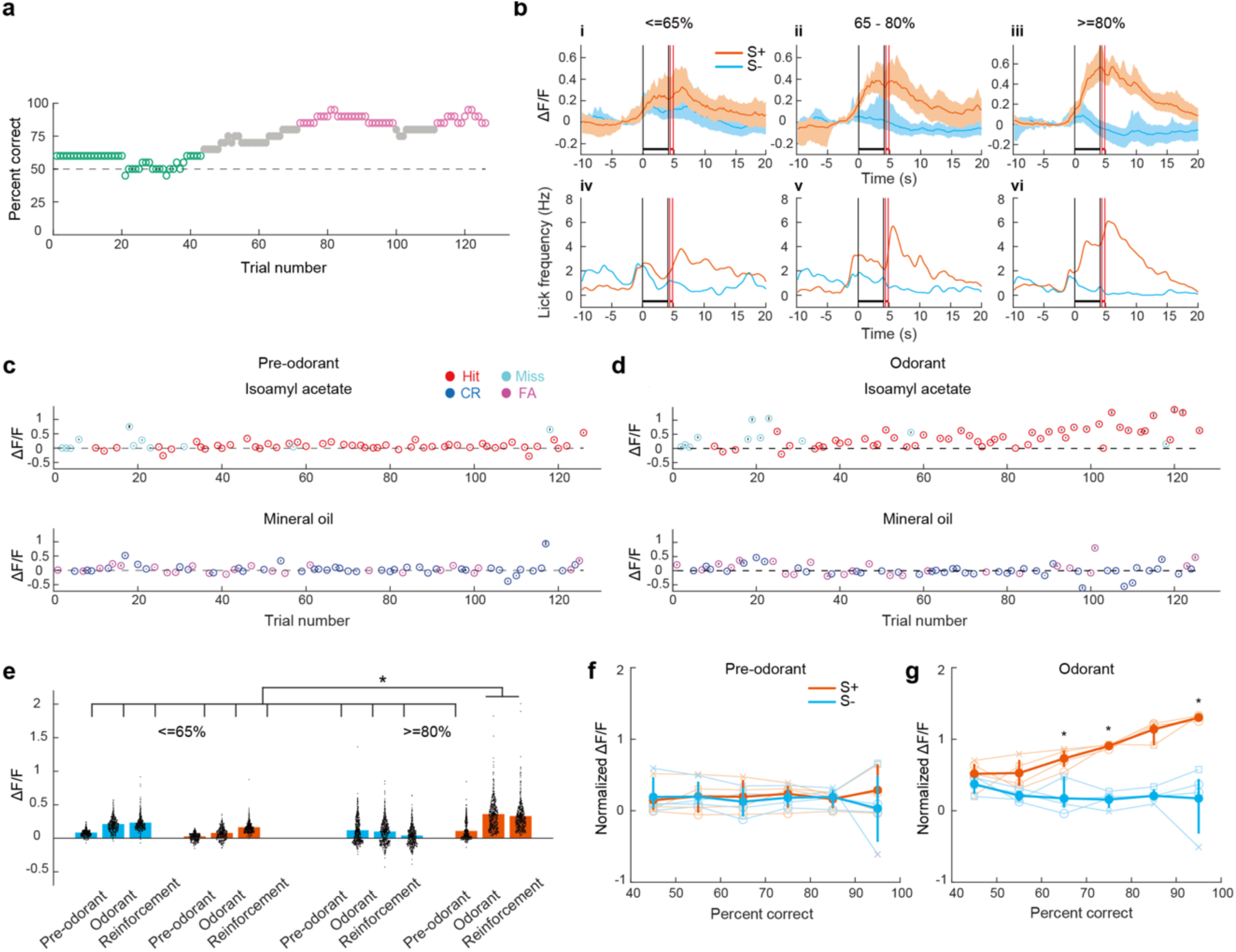
MLIs developed divergent responses during go-no go learning. **a**. Learning curve for a mouse discriminating 1% Iso from MO in the go-no go task. Magenta dots: proficient level (>=80% percent correct), green dots: naïve level (percent correct <=65%). **b**. GCaMP6f fluorescence (ΔF/F) time course averaged over all ROIs and trials falling within different proficiency windows shown for the session whose performance is shown in **a. i**. <= 65% (naïve), **ii**. 65%-80% and **iii**. >= 80% (proficient). Orange: S+, light blue: S-(the shaded area is the 95% CIs). The vertical black lines are odorant onset and removal and the red lines bound the reinforcement period. **c-d**. Per trial ΔF/F averaged over all ROIs for one second before odorant application (**c**, Pre-odorant) and in the last second during odorant application (**d**, Odorant) shown for the session in **a**. Vertical lines around the mean of each trial are the CI. **e**. Violin plot showing per ROI ΔF/Fs for the session shown in **a** for the following time windows: pre-odorant (one second before odorant application), odorant (the last second of odorant application) and reinforcement (one and a half seconds after onset of reinforcement). Per ROI ΔF/Fs are shown for trials falling within different proficiency windows: <= 65% (naïve) and >= 80% (proficient). A GLM indicates that there are significant differences between S+ (orange) and S-(light blue), between time windows and between naïve and proficient mice (p<0.001, 5538 d.f., one mouse, *post-hoc ranksum/t test p value < pFDR=0.043). **f-g**. ΔF/F for four mice averaged over all the ROIs for all trials falling within different proficiency windows. ΔF/F average was calculated for one second before odorant application (**f**) and for the last second during odorant application (**g**). For the data during odorant application (**g**) a generalized linear model (GLM) analysis indicated that there is a statistical significance for the interaction between percent correct and the identity of the odorant (p<0.001, 43 d.f., n=4 sessions, 4 mice, GLM F-statistic=60, p<0.001). GLM did not yield statistically significant differences for pre-odorant data (**f**, p>0.05, 43 d.f., n=4 sessions, 4 mice, GLM F-statistic=0.46, p>0.05). *Post-hoc t-test, p<pFDR=0.018.

### The identity of the stimulus can be decoded from MLI activity in proficient mice

The finding that the MLIs developed divergent responses for S+ vs. S-trials (Fig. 2g) raised the question whether ensemble neural activity encodes for the identity of the stimulus. The learning curve for a session where the mouse achieves >=80% in ∼60 trials is shown in Fig. 3a. The first two principal components for a principal component analysis (PCA) of ΔF/F values for all MLIs in this session showed clear differences between S+ and S-trials for the reward and odorant periods for trials when the mouse was proficient, in contrast to the overlap observed between odorants for the naïve mouse (Fig. 3b). This suggested that odorant identity (S+ vs. S-) could be decoded from ΔF/F values for proficient mice.

**Figure 3.**
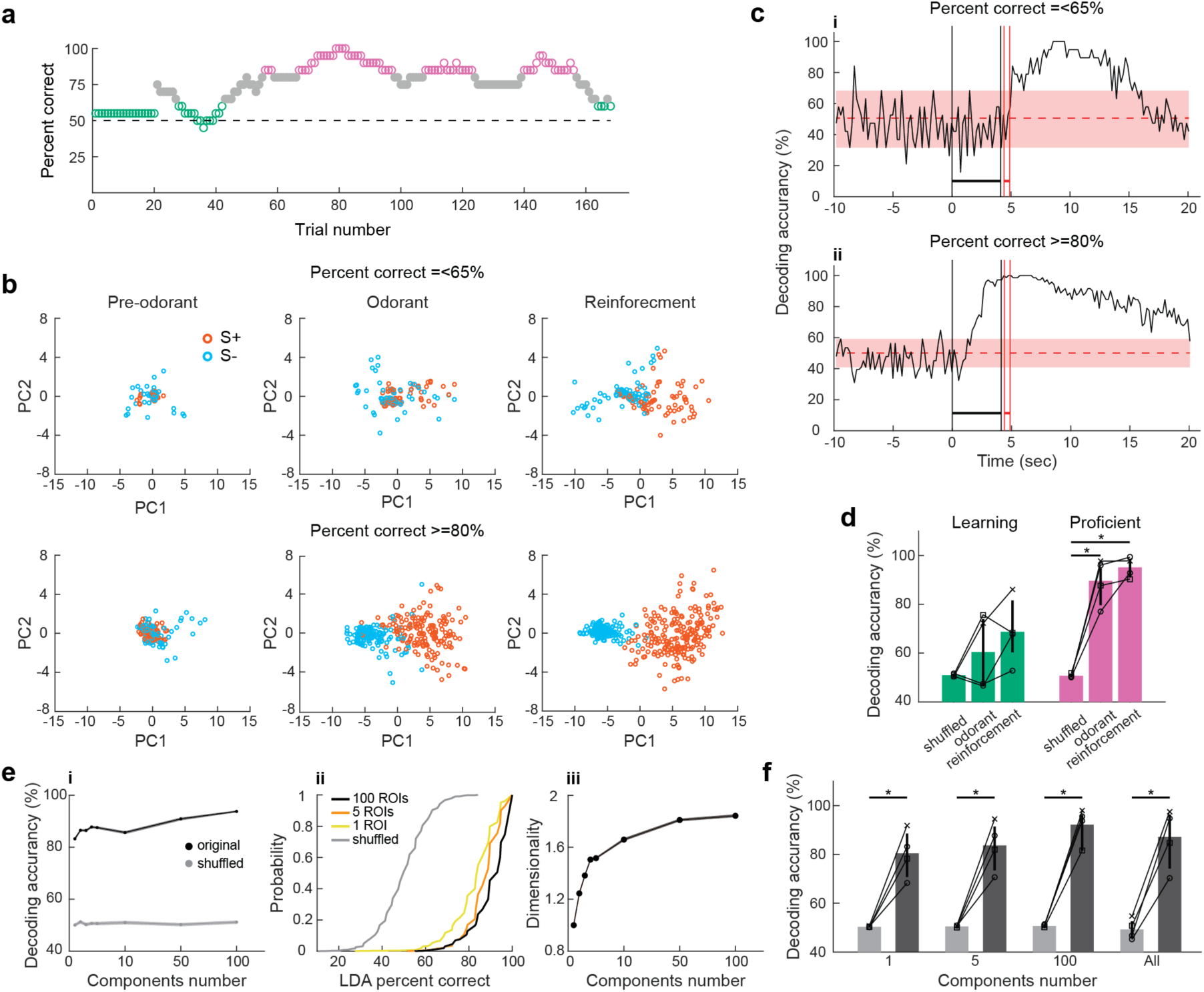
The accuracy of decoding the stimulus increases as the animal learns to discriminate the odorants. Panels **a-c** show behavioral performance (**a)**, PCA analysis (**b)** and LDA analysis (**c)** for one session (one mouse). **a**. Learning curve for a mouse performing the head-fixed go-no go task. Magenta dots: proficient level (>=80% percent correct), green dots: percent correct <=65%. **b**. First two principal components for the PCA for the changes in GCaMP6f fluorescence (ΔF/F) in all the ROIs in the FOV. Top: the mouse was performing =< 65% correct, bottom: the same mouse was performing >=80% correct. The principal components are shown for three time periods: Pre-odorant: 1 sec before odorant addition, odorant: 1 sec before removal of the odorant, and Reinforcement: 1.5 sec after reward. Orange circles: S+ trials, light blue circles: S-trials. **c**. Decoding accuracy for the linear discriminant analysis (LDA) trained to predict odorant identity using ΔF/F for all ROIs in the FOV. Shade: 95% CI for LDA trained with shuffled odorant identity. The vertical black lines are odorant onset and removal and the red lines bound the reinforcement period. **d**. LDA decoding accuracy calculated from the (ΔF/F) for all ROIs in the FOV from data recoded in four mice. A GLM analysis of decoding accuracy yielded statistically significant interactions between (shuffled vs. odor) x (proficient vs. naïve) and for (shuffled vs. reinforcement) x (proficient vs. naïve) (GLM p values <0.01 and <0.05, 24 observations, 18 d.f., n=4 sessions, 4 mice, GLM F-statistic=16.4, p<0.001). The vertical lines show 95% CIs. *Post hoc t-tests corrected for multiple comparison using false discovery rate (FDR, p<pFDR=0.007, n=4 sessions, 4 mice). **e**. Example of LDA decoding analysis performed with subsets of ROIs ranging from 100 to 1 ROI. The analysis was performed for one session for trials where the mouse was proficient (percent correct >=80%). **i**. Decoding accuracy (mean± 95% CI) as a function of the number of ROIs. **ii**. Cumulative probability histograms for the decoding accuracy for the different number of ROIs and for the shuffled LDA. **iii**. Dimensionality as a function of the number of ROIs. **f**. Summary graph showing the decoding accuracy for different numbers of ROIs (mean± 95% CI, 4 sessions, 4 mice). A GLM analysis indicated that the decoding accuracy for the shuffled analysis was statistically different from accuracy with the subsets of ROIs (p<0.001, 72 observations, 54 d.f., n=4 sessions, 4 mice, GLM F-statistic=23.6, p<0.001). *Post-hoc t-tests or ranksum p<pFDR=0.03.

We utilized a linear discriminant analysis (LDA) to determine whether a hyperplane placed in the multidimensional space of ΔF/F values for all MLIs in the ensemble could decode the stimulus. The LDA was trained with ΔF/F for all trials minus one and was queried to identify the odorant in the remaining trial (see Methods). The bootstrapped 95% confidence interval (CI) for an LDA trained after shuffling stimulus was used as a control. The time course for LDA decoding accuracy for the session with the learning curve in Fig. 3a is shown in Fig. 3c (Supplementary Fig. 5a shows the corresponding lick frequency time course). For trials with percent correct <=65%, decoding accuracy raises above the shuffled 95% CI after water reinforcement (Fig. 3ci), while for proficient trials, decoding accuracy starts rising above CI shortly after odorant addition (Fig. 3cii). When tested in four mice LDA decoding accuracy for the odorant and reinforcement periods differed from shuffled LDA (Fig. 3d). A GLM analysis yielded statistically significant interactions between shuffled vs. reinforcement and for the interactions between (shuffled vs. odorant) x (proficient vs. naïve) and (shuffled vs. reinforcement) x (proficient vs. naïve) (GLM p values <0.01 and <0.05, 24 observations, 18 d.f., n=4 sessions, 4 mice, GLM F-statistic 16.4, p<0.001). Post hoc tests corrected for multiple comparison using false discovery rate (FDR)^33^ yielded a statistically significant difference for decoding accuracy for either reinforcement or odorant vs. shuffled for proficient (p<pFDR=0.007, n=4 sessions, 4 mice). Performing the analysis on subsets of ROIs showed that even analysis with a single ROI yields decoding accuracy that is significantly different from the shuffled LDA indicating that the information on the stimulus stored in MLI activity is highly redundant (see Supplementary Note 2, Figs. 3e and f). The LDA analysis revealed that the accuracy for decoding the odorant identity from ΔF/F during the odorant period increases as the animal learns to differentiate between odorants.

### MLI odorant responses switch when stimulus valence is reversed

In order to determine whether the MLIs responded to the chemical identity of the odorant, as opposed to responding to the valence (contextual identity: is the stimulus rewarded?), we reversed odorant reinforcement. When the reward was reversed for a proficient mouse the animal kept licking for the previous rewarded odorant resulting in a fall in percent correct below 50%, and as the animal learned the new valence the percent correct raised back above 80% (Fig. 4a and Supplementary Figs. 6e and 6f). We analyzed how odorant-elicited changes in MLI ΔF/F varied in this reversal task. Fig. 4b shows single trial examples for mean ΔF/F odorant responses when the reward is reversed. MLIs respond to Iso (S+) with an increase in ΔF/F before reversal (e.g. forward trial 52), maintain the odorant increase in ΔF/F to what is now the unrewarded odorant (Iso, S-) immediately following reversal (reverse trial 92) and switch responses with increases to the new rewarded stimulus (MO, S+) when they become proficient in the reversal task (reverse trial 438) (ΔF/F for the last second of the odorant period is shown for all trials in Fig. 4c). When repeated for three different mice the odorant-induced ΔF/F changes reversed for proficient mice (>=80% correct) when the valence was reversed (Fig. 4d). GLM analysis indicates that there is a significant difference in odorant-induced changes in ΔF/F for the odorant and the reversal (p<0.001, 12 observations, 8 d.f., n=3 sessions, 3 mice, GLM F-statistic=14.8, p<0.01). Thus, after successful reversal the ΔF/F time course switched for the two odorants: the stimulus-induced increase in ΔF/F took place for the reinforced odorant, not for the chemical identity of the odorant.

**Figure 4.**
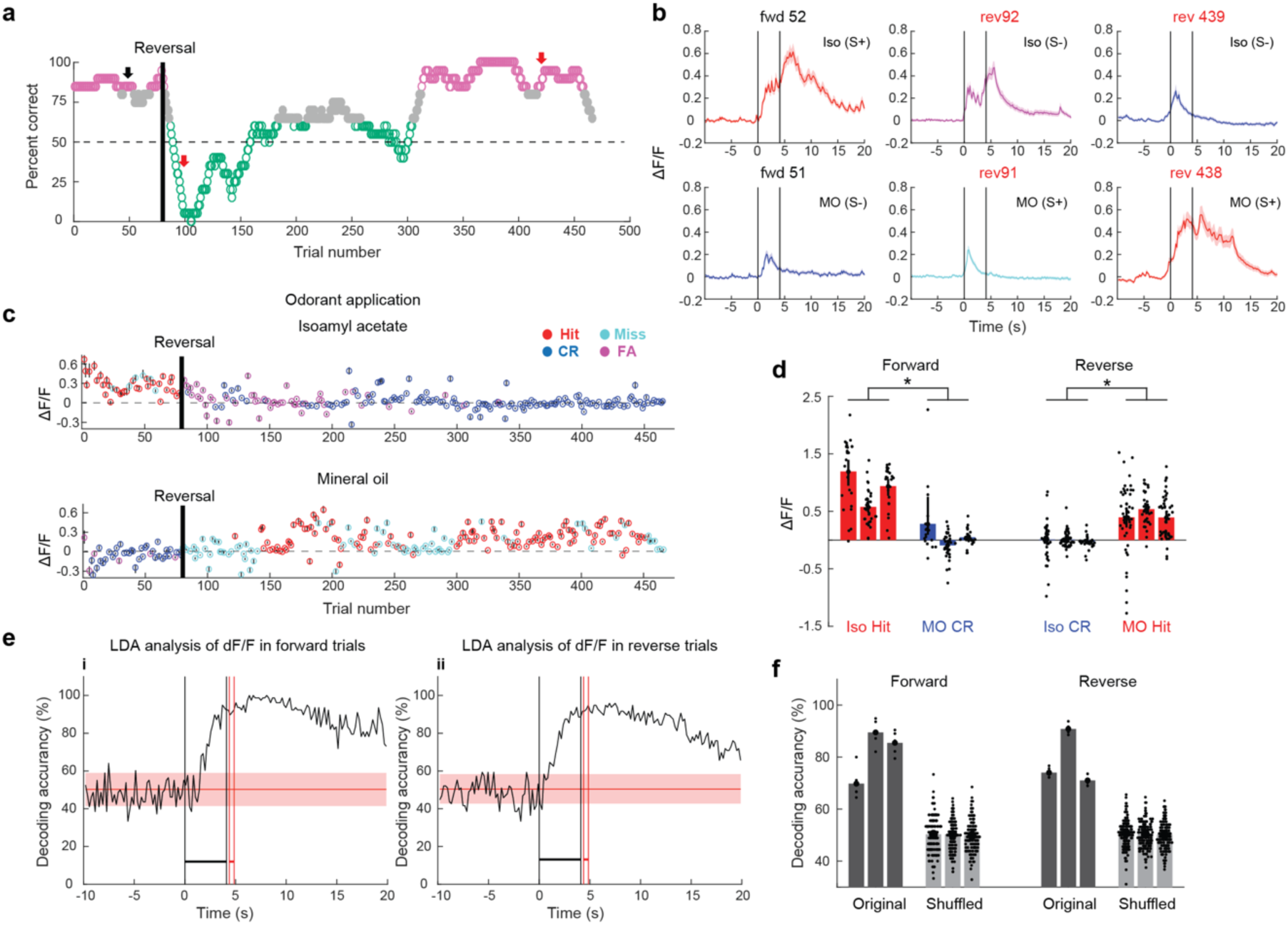
MLI odorant responses switched after reversal. Panels **a-c and e** show behavioral performance (**a)**, ΔF/F (**b** and **c**) and LDA analysis (**e**) for one session where odorant reinforcement was reversed (one mouse). **a**. Learning curve showing the animal’s behavior after reversal. Green <=65%, magenta: >=80% correct. Forward: S+ 1% Iso and S-MO. **b**. Average ΔF/F time courses for example trials before (trials 51 and 52) and shortly after reversal (trials 91 and 92), and after the animal attained proficiency during reversal (trials 438 and 439). The vertical black lines are odorant onset and removal. **c**. Per trial average ΔF/F during the last second of odorant application for the reversal task in Fig. 4a. The upper plot shows the responses to Iso and the lower plot shows responses to MO. **d**. Odorant-induced changes in ΔF/F recorded from three animals computed for trials when the animal was behaving >=80% correct. Vertical lines are 95% CIs. Red: Hit, Blue: CR. A GLM analysis indicates that there is a significant difference in odor-induced changes in ΔF/F for both odorant and reversal (p<0.001, 12 observations, 8 d.f., n=3 sessions, 3 mice, GLM F-statistic=14.8, p<0.01). **e**. Time course for decoding accuracy computed with LDA analysis of ΔF/F data for all ROIs in the FOV. The shade is the 95% CI of decoding accuracy calculated with shuffled odorant identity. The vertical black lines are odorant onset and removal and the red lines bound the reinforcement period. **f**. Decoding accuracy for three animals calculated with odorant-induced changes in ΔF/F from trials where the animal was performing >=80% correct. Vertical lines are 95% CIs. Dark-gray: original, light-gray: shuffled. A GLM analysis indicates that differences in decoding accuracy are statistically significant between original and shuffled (P<0.001), but not between forward and reverse (p>0.05, n=12, 8 d.f., n=3 sessions, 3 mice, GLM F-statistic=16, p<0.001).

Next we computed the accuracy for decoding the reinforced odorant for trials when the animal was proficient in either the forward or reverse trials using LDA analysis. Fig. 4e shows for the reversal session in Fig. 4a that for the proficient animal decoding accuracy rises above the 95% CI calculated with shuffled trials shortly after odorant addition for both forward and reverse trials. Finally, the results of the forward and reverse LDA analysis for three mice that are proficient showed that decoding accuracy for the stimuli differed from shuffled LDA (Fig. 4f). GLM analysis indicates that differences in decoding accuracy are statistically significant between original and shuffled (P<0.001), but not between forward and reverse (p>0.05, 12 observations, 8 d.f., n=3 sessions, 3 mice, GLM F-statistic=16, p<0.001). Finally, decoding accuracy of an LDA analysis for contextual identity for proficient mice did not differ between correct (Hits and CRs), and incorrect trials (Miss and FAs) (Supplementary Note 3, Supplementary Figs. 8,9). These results indicate that for the proficient mouse it is possible to decode contextual identity suggesting that MLI activity encodes for valence.

### Changes in lick rate correlate with changes in MLI activity during reinforcement, but not during odorant application

To explore whether the observed MLI ensemble activity recorded in vermis is directly related to licking, as found in Crus II^34,35^, we examined the correlation between ΔF/F and lick rate. When the animal was proficient the animal licked at least once in each of the two lick segments during application of the S+ odorant, increased licking after receiving the water reward and refrained from licking for the S-trials (Fig. 5a). For the proficient mouse the lick frequency diverges between S+ and S-trials shortly after addition of the odorant (Fig. 5b) and this divergence is evidenced by a large decrease in the p value calculated with a ranksum test comparing licking between S+ and S-trials (Fig. 5c).

**Figure 5.**
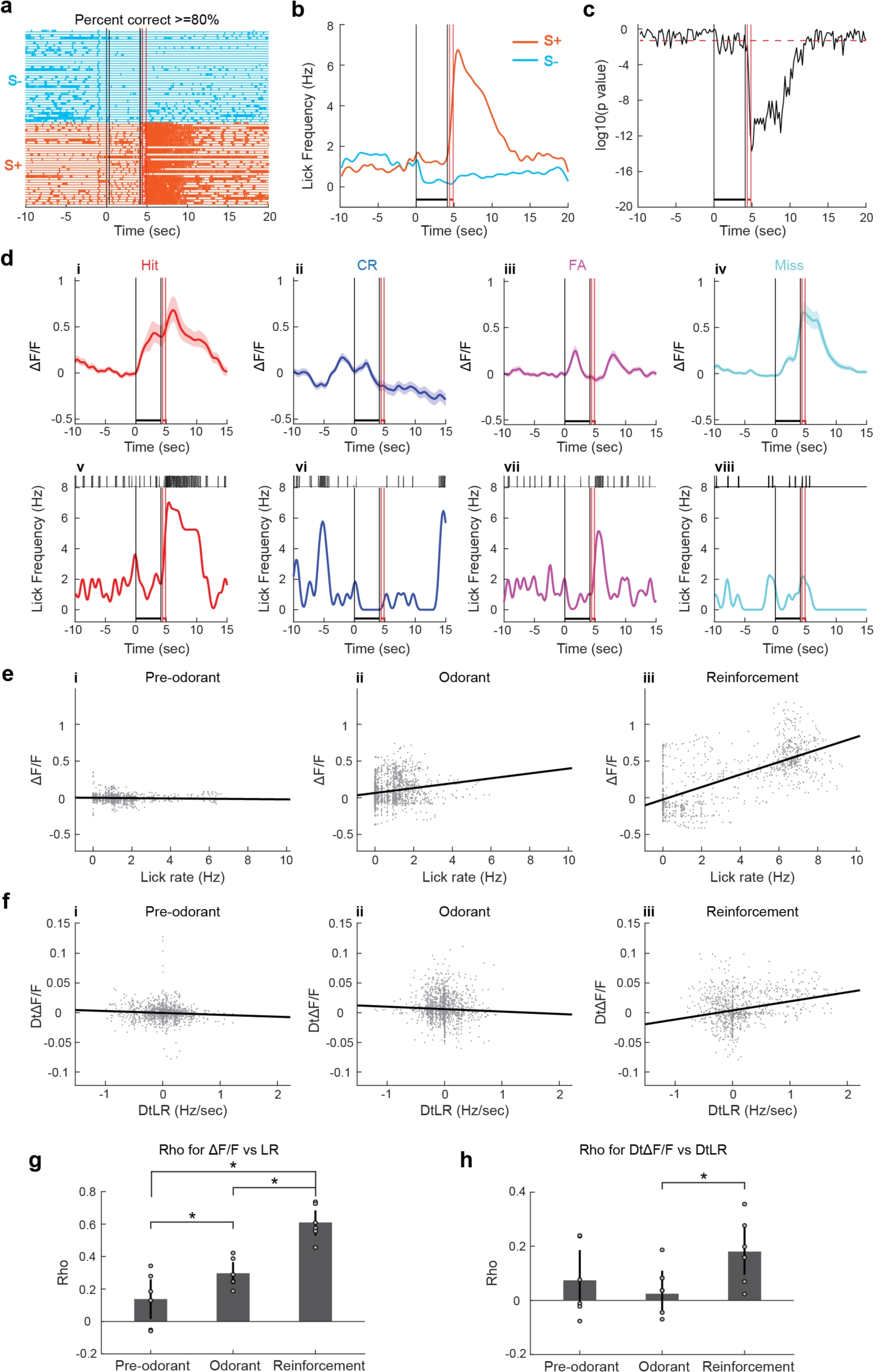
Changes in lick rate correlate with changes in MLI activity during the reinforcement period, but not during the odorant period. Panels **a-d** are examples of lick traces (**a**), lick frequency (**b**), lick ranksum p value (**c**) and ΔF/F (**d**) for one session (one mouse). **a**. Examples of per trial lick traces when the mouse was learning to discriminate the odorants when the animal was proficient (>=80% correct, orange traces: S+ trials, light blue traces: S-trials). **b**. Average lick frequency for S+ (orange) and S-(light blue) trials shown in **a**. **c**. p value for a ranksum test estimating the difference in licks between the S+ and S-odorants for the example in **a**. **d**. Single trial examples of average ΔF/F (±CI, shade, n=105 ROIs) (**i-iv**) and the corresponding lick frequency and lick traces (**v-viii**). **e**. Relationship between the per trial average ΔF/F and the lick frequency shown per time point for all trials in a go-no go session (one session, one mouse). Time points are segregated within the last second before odorant addition (Pre-odorant), during the last second of odorant addition (Odorant) and during the 1.5 seconds after reward (Reinforcement). Correlation coefficients and p values for these time periods are: Pre-odorant: 0.058, p<0.05. Odorant: 0.19, p<0.001. Reinforcement: 0.73, p<0.001. *p<pFDR=0.05, n=6 sessions, 5 mice. **f**. Relationship between the derivative of average ΔF/F and the derivative of lick frequency shown per time point for all trials in a go-no go session (one session, one mouse). Time points are segregated within the last second before odorant addition (Pre-odorant), during the last second of odorant addition (Odorant) and during 1.5 seconds after reward (Reinforcement). Correlation coefficients and p values for these time periods are: Pre-odorant: −0.076, p<0.01. Odorant: −0.045, p>0.05. Reinforcement: 0.28, p<0.001. *p<pFDR=0.016, n=6 sessions, 5 mice. **g**. Correlation coefficients for the relationship between the average ΔF/F and lick frequency for six sessions (five mice). The correlation coefficient is significantly different between reinforcement and odorant (*t test p<pFDR=0.05, n=6 sessions, 5 mice). **h**. Correlation coefficients for the relationship between the derivative of average ΔF/F and the derivative of lick frequency for six sessions (five mice). The correlation coefficient is significantly different between reinforcement and odorant (*t test p<pFDR=0.016, n=6 sessions, 5 mice). In b, c and d the vertical black lines are odorant onset and removal and the vertical red lines bound the reinforcement period. All data shown in this figure are for proficient mice (percent correct>=80%).

In order to explore the relationship of MLI activity to licking we proceeded to examine the correlation between ΔF/F time course and the lick rate (LR) and between the derivative of the ΔF/F time course (D_t_ΔF/F) and the derivative of the lick rate (D_t_LR)^34^ and plotted their relationship during different time periods for proficient mice. Fig. 5d shows examples for several trial outcomes for the time course for ΔF/F and the lick rate. Examples of these correlations for a single session for a proficient mouse are shown in Figs. 5e and f. In this example the correlation between ΔF/F and LR was larger during the reinforcement period (ρ=0.73, p-value<0.001) compared to the odorant (ρ=0.19, p-value<0.001) and pre-odorant (ρ=-0.06, p-value<0.05) periods (Fig. 5e). Similarly, the correlation between D_t_ΔF/F and D_t_LR is significant for the reinforcement period (ρ=0.28, p<0.001), and is smaller for both the pre-odorant (ρ=-0.08, p<0.05) and odorant (ρ=-0.045, p>0.05) periods (Fig. 5f). We proceeded to compare the correlations between these parameters in several mice. As found in Crus II, ΔF/F and LR (Fig. 5g) and D_t_ΔF/F and D_t_LR (Fig. 5h) are positively correlated during the reinforcement period. Interestingly, the correlations are lower during the pre-odorant and odorant periods (Figs. 5g,h). A t test indicated that there is a statistically significant difference in the correlation coefficient between the reinforced vs. the odorant or pre-odorant periods for ΔF/F vs. LR (p<pFDR=0.05, n=6 sessions, 5 mice) and between the reinforced and odorant periods for D_t_ΔF/F vs. D_t_LR (p<pFDR=0.016, n=6 sessions, 5 mice). These data suggest that changes in ΔF/F during the reinforcement period reflect changes in lick activity, while changes in ΔF/F during the odorant period are less dependent on licks, and maybe dependent on multiple variables. We performed complementary studies of ΔF/F in CR trials when the mouse did not lick and for time-courses aligned to the beginning of ΔF/F changes after odorant addition (Supplementary Note 4 and Supplementary Figs. 10 and 11). Under these conditions, ΔF/F and lick rate are uncorrelated, consistent with the potential lack of direct relationship between changes in ΔF/F and lick rate during the odorant period.

### Do the MLIs respond to reward value?

The correlation between ΔF/F time course and the lick rate (Fig. 5e) raises the question whether ΔF/F reflects the reward value as opposed to the valence of the odorant. Here we define the valence of the odorant as indicating whether the stimulus is good or bad^36^. Thus, valence is a binary measure of an emotion that is reflected by the motivation to receive reward, whereas reward value, in the present paradigm, is related to the amount of sugar water delivered. To evaluate whether ΔF/F changes are dependent on reward value we recorded changes in ΔF/F from mice performing a go-go task where both odorants were rewarded equally and we varied the volume of sugar water delivered for successful trials. As expected, the mouse responded to both odorants (Supplementary Fig. 12a). Also, as expected, when the volume of reward was increased, the lick frequency increased during the delivery of sugar water (wet licks, Supplementary Fig. 12b), but not during dry licking before reward (dry licks, Supplementary Fig. 12b). A GLM analysis yielded a statistically significant difference for dry vs. wet licking and for the interaction between the volume of sugar water delivered and dry vs. wet (p<0.001, 198 observations, 194 d.f., 1 session, 1 mouse, GLM F-statistic=21.8, p<0.001). In contrast, a GLM analysis of the changes in ΔF/F as a function of volume delivered in dry and wet lick conditions did not yield statistically significant changes (Supplementary Fig. 12c, p>0.05, 198 observations, 194 d.f., 1 session, 1 mouse, GLM F-statistic=1.7, p>0.05). Finally, if MLI activity reflected reinforcement value ΔF/F should show a positive correlation with volume of sugar water delivered (see Fig. 3 of ^34^). We did not find a significant correlation between lick frequency and ΔF/F for either dry (Supplementary Fig. 12di, ρ=0.16, p>0.05) or wet licking (Supplementary Fig. 12dii, ρ=-0.66, p>0.05). This experiment indicates that MLI activity does not reflect reward value, and is consistent with MLI activity reflecting valence.

### The main contributor during odorant application to a GLM model fit to MLI activity is the contextual identity of the odorant

In order to understand the contribution of different variables to MLI activity we setup a GLM to quantify the dependence of the average ΔF/F on the different behavioral and stimulus variables^37^. We included “event” variables, “whole trial” variables and “continuous” variables (Fig. 6a, Methods). Continuous variables quantified kinematics including lick rate and the derivative of lick rate (Fig. 5) and body velocity and body acceleration of movements made by the animal during the trial (Supplementary Fig. 3). The identity of the odorant (S+/S-odorant) was an event variable that increased from zero to one during the time for odorant application. Finally, whole trial variables were accuracy (1 for correct and 0 for incorrect responses), reinforcement history (1 for reinforcement in the last trial, 0 otherwise) and percent correct behavior calculated in a window of 20 trials.

**Figure 6.**
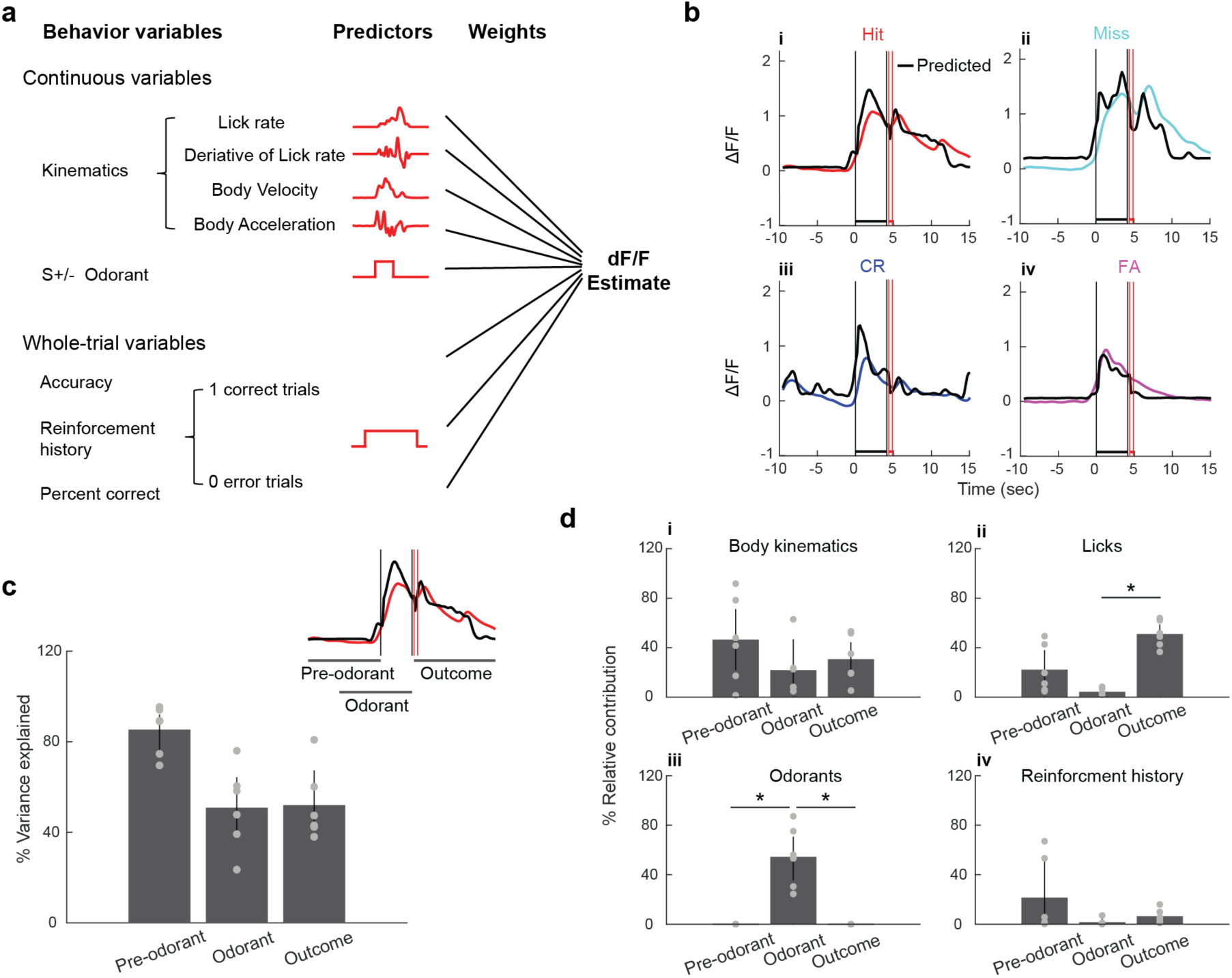
Fit of MLI activity with a generalized linear model. **a**. Schematic illustration of the variables used by the GLM model to quantify the relationship between average ΔF/F and behavioral variables and odorant valence on a per trial basis. **b**. Examples of GLM best-fit (black) and actual ΔF/F Ca^2+^ traces for Hit (**i**, red), Miss (**ii**, cyan), CR (**iii**, blue) and FA (**iv**, magenta) trials. The vertical black lines are odorant onset and removal and the red lines bound the reinforcement period. **c**. Percent of the variance explained by GLM in the different time periods (Pre-odorant, Odorant and Outcome). The inset shows the time intervals used for the different periods. **d**. Relative contribution of kinematics (**i)**, licks (**ii**), odorants (**iii**) and reinforcement history (**iv**) to the GLM fit for fits in the different time periods (pre-odorant, odorant, outcome). The results showed that odorants and licks play an important role in contributing to the activity during odorants and outcome periods respectively, while kinematics contribute to the activity in all three time periods. Asterisks for Licks (**ii**) and Odorants (**iii**) denote significant differences determined by ranksum and t test (p<pFDR=0.033, n=6 sessions, 5 mice). There are no significant differences for Kinematics (**i**) or Reinforcement history (**iv**) (p>pFDR=0.017, ranksum test).

Black traces in Fig. 6b show examples of the fit of the GLM model to average ΔF/F time courses for four example trials with different outcomes. The fit traces in black largely overlap with the recorded data. When quantified over the different periods within a trial (pre-odorant, odorant and reinforcement) we found that GLM explained a substantial percent of the average ΔF/F variance ranging from 23.5% to 95% (Fig. 6c). We next quantified the relative contribution of the different variables to the ensemble activity in the three time periods. Fig. 6d shows the contributions to the GLM fit by the variables that make the largest percent contribution to the fit (body kinematics, licks, reinforcement history and odorants). Odorant (S+ vs. S-) is the dominant contributor to the activity during odorant application (Fig. 6d,iii, asterisks show differences with ranksum or t test p<pFDR=0.033, n=6 sessions, 5 mice). Furthermore, variables describing the licks contribute to the GLM fit during the outcome period (Fig. 6d,ii, * p<pFDR=0.016 for a t test, n=6 sessions, 5 mice). In contrast, the other two variables (body kinematics and reinforcement history) do not differ in contribution during the different periods (p>pFDR=0.017, ranksum test). These data indicate that odorant identity contributes to modeling MLI activity during the odorant application period while licks contribute to MLI activity during the reinforcement period.

### Chemogenetic inhibition of MLI activity leads to impaired go-no go learning

In order to determine whether activity of MLIs plays a role in behavioral responses in the go-no go task we used a Cre-dependent AAV virus to express the inhibitory DREADDs receptor hM4Di in MLIs in six PV-Cre mice (hM4Di group)^38^. To control for off-target effects of clozapine-N-oxide (CNO)^39^ we injected PV-Cre mice with Cre-dependent mCherry AAV virus in another group of six mice (control group) (see Supplementary Fig. 13a,b for virus expression). One to two weeks later animals from both groups were trained to differentiate two odorant mixtures (S+: 0.1% of 60% heptanal+40% ethyl butyrate, S-: 0.1% of 40% heptanal+60% ethyl butyrate) for 3-4 sessions encompassing 400-500 trials in the go-no go task. The animals were injected intraperitoneally (IP) with saline 40 minutes before the start of the sessions (control-saline and hM4Di-saline). One to two weeks later mice were trained again to differentiate between the same odorant mixtures, but they were injected IP with CNO (3mg/kg) 40 minutes before the start of the session (control-CNO and hM4Di-CNO).

Figs. 7a,i and b,i show representative examples of the behavioral performance for the four conditions. Mice in all groups with the exception of the hM4Di-CNO attained proficiency (>=80% percent correct) (Figs. 7a,ii and 7b,ii). GLM analysis indicates that there is a statistically significant interaction between CNO drug treatment and hM4Di expression (p<0.01, 24 observations, 20 d.f., n=6 mice, GLM F-statistic=13.7, p<0.001) and post hoc tests indicate that the hM4Di expressing group differs between CNO and saline (p<pFDR=0.025, n=6 mice per group), while there are no significant differences between CNO and saline for control mice (p>pFDR, n=6 mice per group), indicating an effect of CNO-induced inhibition of MLIs expressing hM4Di on behavioral output and the absence of off-target CNO effects.

**Figure 7.**
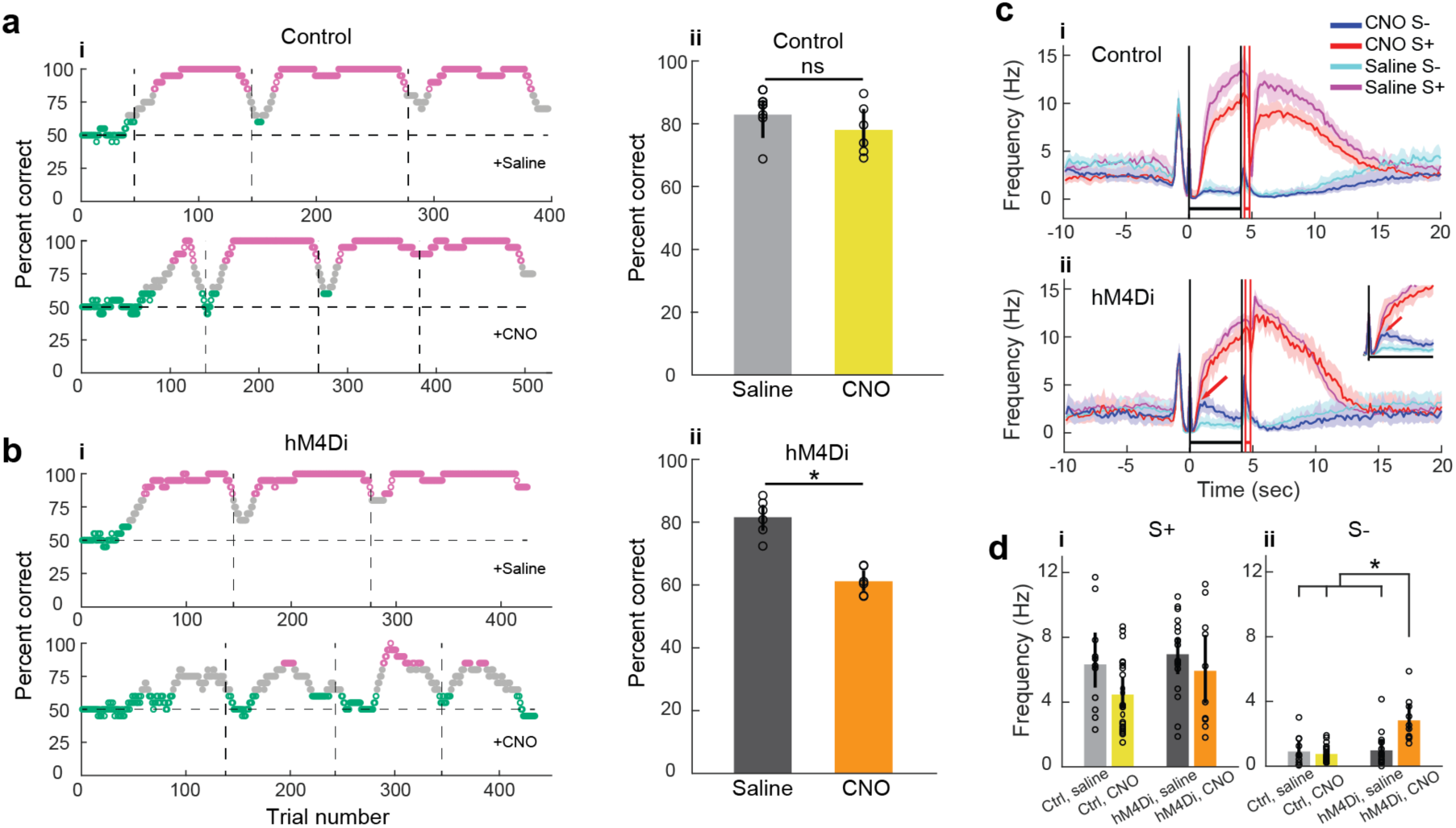
Chemogenetic inhibition of MLI activity impairs associative learning. **a. and b. a,i and b,i**. Examples of behavioral performance in a go-no go task where mice learned to differentiate between two odorant mixtures (S+: 0.1% of 60% heptanal+40% ethyl butyrate, S-: 0.1% of 40% heptanal+60% ethyl butyrate). Green: <=65% correct, magenta: >=80% correct. The vertical lines are the boundaries between different daily sessions. **a,ii and b,ii**. Behavioral performance (mean± 95% CIs, n=6). **a**. Mice expressing mCherry in MLIs (control mice). **b**. Mice expressing hM4Di in MLIs. A GLM analysis indicates that there is a statistically significant interaction between CNO drug treatment and hM4Di expression (p<0.01, 24 observations, 20 d.f., n=6 mice, GLM F-statistic=13.7, p<0.001) and post hoc unpaired t tests indicate that the hM4Di expressing group differs between CNO and saline (asterisks, p<pFDR=0.025, n=12 mice). **c**. Average lick frequency time course for the control (**i**) and hM4Di (**ii**) groups (mean± 95% CI, shade). The arrow in the inset shows that for S-trials of CNO treated hM4Di animals (blue), the animal kept licking for a longer time than S-trials for mice injected with saline (cyan). The vertical black lines are odorant onset and removal and the red lines bound the reinforcement period. **d**. Mean lick frequency (±CI) during the initial portion of the odorant application period (0.8 to 1.8 sec) for mice performing >=80% correct (**i**: S+, **ii**: S-). GLM analysis did not find a statistically significant difference for treatment or hM4Di expression (or interactions) for S+ (p>0.05, 60 observations, 56 d.f., n=6 mice, GLM F-statistic=2.96, p>0.05), but found a difference for CNO treatment x hM4Di expression for S-(p<0.001, 60 observations, 56 d.f., n=6 mice, GLM F-statistic=12.8, p<0.001). *p<pFDR=0.03 for post-hoc t-test.

We proceeded to compare the time course for lick frequency between different conditions for trials when the mouse was proficient (>=80% correct, Fig. 7c). Interestingly, the time course for lick frequency differs between hM4Di mice injected with CNO and the other conditions (Figs. 7c,i and 7c,ii). With the exception of hM4Di + CNO, shortly after odorant addition (∼0.8 sec) the lick frequency for S-does not increase beyond ∼1 Hz while the lick frequency for S+ keeps increasing beyond 8 Hz (Fig. 7c). In contrast, for hM4Di + CNO lick frequency for S-keeps increasing beyond 1 Hz and does not diverge from the S+ time course until it reaches 2.5 Hz at a later time point (∼1 sec, Fig. 7c,ii) likely reflecting slow decision-making. This would explain why the hM4Di + CNO mice accumulated errors in the go-no go task (Fig. 7bii). In order to quantify this difference in lick frequency we calculated the lick frequency 0.8 to 1.8 seconds after odorant addition (Fig. 7d). For S+ GLM analysis did not find a statistically significant difference for treatment or hM4Di expression (or interactions) for the rewarded odorant trials (Fig. 7di, p>0.05, 60 observations, 56 d.f., n=6 mice, GLM F-statistic=2.96, p>0.05). Yet GLM found a statistically significant difference for S-for the interaction between CNO and hM4Di expression (Fig. 7dii, p<0.001, 60 observations, 56 d.f., n=6 mice, GLM F-statistic=12.8, p<0.001). We obtained a similar impairment of performance in a separate set of experiments with two hM4Di mice and two controls where animals discriminated between 1% Iso and MO, and CNO was applied when the animal was naïve, and we reversed the reward (Supplementary Fig. 13). These data indicate that inhibition of MLIs causes a slower differential lick response to odorants leading to impaired behavioral performance.

## Discussion

We found that vermal MLIs developed a differential response to odorants in the go-no go task that switched when the valence was reversed. Decoding analysis revealed that when the animal was proficient the contextual identity of the odorant could be decoded from MLI responses. GLM analysis revealed that contextual identity made a large contribution to the fit of MLI activity during the odorant application period. Chemogenetic inhibition of MLIs impaired achievement of proficient discrimination of odorants. These data indicate that MLIs play a role in associative learning by encoding valence.

The cerebellum has been implicated in mediating supervised learning through an iterative process whereby the response to an input is evaluated against a desired outcome, and errors are used to adjust adaptive elements within the system^8-10,40^. CFs carrying error signals make profuse synaptic connections on the dendrites of PCs and elicit powerful excitatory dendritic Ca^2+^ spikelets^41-44^. Furthermore, CFs also signal reward prediction^11,12^ or decision-making errors^13^, and the cerebellum modulates association pathways in VTA enabling a cerebellar contribution to reward-based learning and social behavior^14^. The increase in Ca^2+^ mediates LTD in subsets of synapses innervated by co-activated GC parallel fibers (PFs) carrying sensorimotor information relevant to learning^45,46^. However, recent studies by Rowan et al.^25^ revealed that increasing feed forward inhibition by MLIs can switch the valence of plasticity from LTD to LTP (also, see^47^). In addition, adaptive changes in the vestibulo-ocular reflex elicited by CF optogenetic activation switched from increase to decrease depending on whether MLIs were co-activated^25^. Finally, MLIs gate supralinear CF-evoked Ca2+ signaling in the PC dendrite^48^. These studies suggest that the valence of learning is graded by MLI activity.

Here we provide evidence for the involvement of MLIs in conveying information on contextual identity of a stimulus in associative learning. We do not find that the MLIs respond to odorants per se. Rather, the reversal experiment (Fig. 4) and the similar ΔF/F responses and stimulus decoding for correct and incorrect behavioral response trials (Supplementary Fig. 8) indicate that MLIs respond to contextual odorant identity: “is this the rewarded odorant?”, which is directly related to valence, a binary measure of an emotion reflected by the motivation to receive reward^36^. Furthermore, our results in the go-go task where both odorants are rewarded with varying volumes of sugar water (Supplementary Fig. 12) are consistent with the response reflecting valence (as opposed to value). Thus, we postulate that the MLI response during the odorant period is related to valence that reflects reward expectation, consistent with the fact that GCs in lobe VI were found to respond to reward expectation^29,30^.

A question that arises is which circuit mechanism is responsible for the decreased behavioral performance after chemogenetic inhibition of MLI activity (Fig. 7 and Supplementary Fig. 13). MLIs receive sensorimotor information from multiple GCs through PF input and *in vivo* studies have found remodeling of MLI receptive fields upon repeated electrical stimulation of the skin^49^. In addition, plasticity in PF-MLI synapses are postulated to increase the information capacity of the MLI-PC network and richness of PC output dynamics^10,16^, and a model of PF-MLI plasticity has been proposed^50^. If long term plasticity of PF-MLI synapses is responsible for the large change of MLI responsiveness found here upon reversal of stimulus valence it is likely that the error signal would be provided by CF spillover resulting in highly redundant stimulation of stellate cells (SCs)^51,52^. We developed a model of the MLI/PC circuit described in Supplementary Note 5 (Supplementary Figs. 14,15) that suggests that plasticity in SC-PC synapses^53^, PF-SC synapses^50^, complemented with CF spillover acting on the feedforward disinhibitory MLI circuit described recently by Arlt and Hausser^52^, complemented with plasticity in SC-PC synapses^53^, PF-SC synapses^50^, would explain the changes in behavior we find after chemogenetic inhibition of MLI activity. Finally, we found that the divergence between the time courses for lick frequency between S+ and S-took place at a later time when MLIs were inhibited by chemogenetics (Fig. 7c,ii, Supplementary Fig. 13g) likely reflecting slow decision-making, consistent with a role for the CF/PC circuit in reward timing prediction^54^. Future studies are necessary to understand the role of plasticity in the PF-MLI-PC circuit in associative learning.

Interestingly, odorant responses have been reported in the cerebellum. Studies in sexually trained male rats found that female bedding or almond smell elicited increased cFos immunoreactive GCs in the vermis compared to rats exposed to clean air^55^. Furthermore, sexual experience increased the number of cFos positive GCs. In humans odorants induced significant activation of the cerebellum^56^. In addition, human cerebellar lesions caused olfactory impairments in the contralesional nostril, and elicited sniffs with lower overall airflow velocity compared to controls^57^. These findings implicate an olfactocerebellar pathway prominent in odorant identification and detection that functionally connects each nostril primarily to the contralateral cerebellum. Our study provides further evidence for involvement of the cerebellum in olfactory tasks.

Here we found changes in MLI activity during learning in the go-no go associative learning task in lobule VI where GCs^29,30^ and CF^58^ activity was proposed to encode aspects of reward signaling. Recent work on the contribution of cerebellar processing to execution of reward-driven behaviors indicates that multiple cerebellar regions are involved, including central and lateral cerebellum^11,29,30,58,59^. As behavioral tests are refined, it is likely that differences in how these regions process reward will emerge, as suggested by the recent work on climbing-fiber signaling^11^. Furthermore, we focused our recordings on the more superficial regions of the molecular layer and therefore, most of our measurements are from SCs. Recordings from synaptically connected pairs of MLIs and PCs in slices showed that mean the amplitude of synaptic currents decreases with distance from the PC layer, suggesting a stronger impact of basket versus SC inhibition on PC firing^60^. Recent recordings of PC spikes in vivo following genetic deletion of MLIs confirm this prediction^47^ which is in accord with the morphological diversity of MLIs^61^. Finally, the effect of chemogenetics (Fig. 7) should be on both stellate and basket cells, and future experiments are necessary to differentiate between the roles of the two cell types.

Our findings show that differential MLI activity develops during learning in an associative learning task and that MLI activity switches when the rewarded odorant is reversed. We find that inhibition of MLI activity elicits decreased behavioral performance in the go-no go task. Our data indicate that MLIs have a role in learning valence. This would likely increase the information capacity of the MLI-PC network and richness of PC output dynamics^10,62^.

## Acknowledgments

We thank Ms. Nicole Arevalo for animal husbandry, Ms. Dnate’ Baxter for laboratory support, Ms. Arianna Gentile-Polese for help with the summary figure and Mr. Jesse Gilmer with help coding dimensionality. We thank Mr. Matt I. Becker, Mr. Jesse Gilmer and Dr. Abigail Person for discussions. This research was supported by NIDCD R01 DC000566, NINDS U01 NS099577, NSF CBET-1631912, NSF BIO-1926676 and ANR-18-CE16-0010-01.

## Author contributions

M.M., I.Ll. and D.R. designed all experiments aided with discussions with E.G., B.O., F.S. and G.F., while M.M. G.F. and D.R. performed them. E.G., G.F. and B.O. designed, setup and maintained the custom two-photon microscope, M.M. and D.R. set up the olfactometer. D.R. wrote all programs for data collection and analysis. F.S. performed circuit modeling, M.M. and D.R. analyzed the data. M.M. and D.R. generated the figures. All authors discussed the results. M.M. and D.R. wrote the manuscript, and all authors participated in commentary and revision of the manuscript.

## Methods

### Animals

All animal procedures were performed in accordance with protocols approved by the Institutional Animal Care and Use Committee of the University of Colorado Anschutz Medical Campus. Mice were bred in the animal facility. We used both male and female adult Parvalbumin-Cre (PV-Cre, Stock number 008069, Jackson Laboratory, USA) mice and wild-type C57BL/6J mice. The animals were housed in a vivarium with a 14/10h light/dark cycle. Food was available ad libitum. Access to water was restricted in for the behavioral training sessions according to approved protocols, all mice were weighed daily and received sufficient water during behavioral training to maintain >=80% of original body weight.

### Immunohistochemistry

To perform immunostaining, mice were sacrificed and transcardially perfused with ice cold 4% paraformaldehyde (Electron Microscopy Sciences, USA), followed by incubation in 30% sucrose (Sigma-Aldrich, USA). After the brain was incubated in the sucrose solution 60 μm thick slices were cut with a cryostat. The slices were imaged using a confocal laser scanning microscope (Leica TCS SP5II, Germany or Nikon A1R, Japan) to determine the GCaMP expression patterns in the cerebellum. The slices were counterstained with DAPI (Thermo Fisher Scientific, USA).

### Window implantation

Adult mice (8 weeks or older) were first exposed to isoflurane (2.5%) and then maintained anesthetized by intraperitoneal ketamine-xylazine injection (100 μg/g and 20 μg/g). A craniotomy was made over the vermis of cerebellum centered at midline 6.8 mm posterior to Bregma leaving the dura intact (lobule VI). A square glass window (2 mm x 2 mm) of No 1 cover glass (0.13 to 0.17 mm thick, Thermo Fisher Scientific, USA) was placed over the craniotomy and the edges were sealed with cyanoacrylate glue (3M, USA). The window was further secured with Metabond (Parkell, USA), and a custom-made steel head bracket was glued to the skull.

### Virus expression of GCaMP

In order to express GCaMP6f in cerebellar MLIs we injected the vermis of the cerebellum in three adult PV-Cre mice with 2.0 μl of AAV1-Syn-Flex-GCaMP6f (Addgene, USA) (6.8 mm posterior to Bregma, bilaterally ±0.5 mm lateral to midline and 200-400 μm below the brain surface)^63^. The viral infection method we use has been reported to express GCaMP in MLIs, and not in PCs^64,65^. Supplementary Fig. 2 shows that indeed expression of GCaMP6 takes place only in MLIs. Note the absence of fluorescence from GCs as well as from PC somata and dendrites. Expression in MLIs may reflect the differential activity of the hSyn promoter, and has been described for a different genetically-encoded Ca^2+^ indicator^65^. In one C57BL/6 animal each we used AAV5-Syn-GCaMP6s or AAVrg-Syn-jGCaMP7f (Addgene, USA) with similar results to those obtained with GCaMP6f (Supplementary Fig. 16a) and therefore the data were pooled. After the injection, the animals were maintained for at least three weeks before behavior training and imaging.

### Go-No Go training

Mice were water deprived by restricting daily consumption of water to 1-1.5 ml. Mice were monitored for signs of dehydration or a decrease in body weight below 80% of the initial weight. If either condition occurred the animals received water ad-lib until they recovered. When the animals were thirsty, they were trained in a head-fixed olfactory go-no go task with (1% Iso vs MO odorant application, Sigma-Aldrich, USA)^26,66^. Licks were monitored by an electrical circuit monitoring the resistance between the lick spout and the floor in an olfactometer that controlled valves to deliver a 1:40 dilution of odorant at a rate of 2 lt/min. The water-deprived mice started the trial by licking on the water port. The odorant was delivered after a random time interval ranging from 1 to 1.5 seconds. In S+ trials, the mice needed to lick at least once in two 2 sec lick segments to obtain a reward (0.1 g/ml sucrose water) (Fig. 1a). The inter trial interval (ITI) was 22.3-22.8 sec. In S-trials, the mice need to refrain licking one of the two 2 sec segments to avoid a longer inter-trial interval (22.3-22.8 + 10 sec). The animal’s behavior performance was evaluated in a sliding window of 20 trials and the calculated value was assigned to the last trial in the window. Therefore, it estimates the performance in the last 20 trials. The percent correct value represents the percent of trials in which the animal successfully performed appropriate actions, and we considered the animal proficient if percent correct performance is above 80%. In reverse go-no go training sessions, the rewarded and un-rewarded odorants were switched. Movement of the mouse was imaged in the infrared to prevent light interference with the non-descanned detection in the visible using a 1 Megapixel NIR security camera (ELP-USB100W05MT-DL36, Amazon.com, USA) at 30 frames per second. Velocity of body movement was estimated using the Farneback algorithm coded in Matlab^67^ (Mathworks, USA). Measurement of body movement with a single camera gives limited information. Five mice were used for the go-no go experiments. Four mice were imaged when they were naïve. The window became opaque for two of the mice preventing MLI imaging for the reversal experiment.

### Go-Go training

For the experiment in Supplementary Figure 12 both odorants were rewarded with the same volume of sugar water, and the volume of sugar water reward was varied. The two odorants in this experiment were either 1% Iso and MO, or the same odorant 1% Iso and 1% Iso. The results were similar for both odorant pairs, and the analysis was performed for all trials regardless of odorant pair.

### Behavioral performance recorded after chemogenetic inhibition of MLI activity

For chemogenetic inhibition of MLI activity^38,39^, 1.2 ul of AAV8-hSyn-DIO-hM4D(Gi)-mCherry virus (Addgene, USA) were bilaterally injected into 6 PV-Cre animals bilaterally ±0.5 mm lateral to midline, 6.8 mm posterior to Bregma and 200-400 μm below the brain surface. For control an AAV8-hSyn-DIO-mCherry virus (Addgene, USA) was injected in the same position in 6 PV-Cre animals. This viral infection method results in expression of the protein in MLIs, and not in PCs^64,65^. We performed two separate experiments: 1) For the experiments in Fig. 7 four weeks after injection the animals were trained to proficiency in the go-no go task with 1% Iso (S+) vs MO (S-). When they reached a proficient level, the animals rested for one to two weeks, and were then injected intraperitoneally (IP) with saline 40 minutes before starting the session and were trained to discriminate between two odorant mixtures: 0.1% of 60% heptanal+40% ethyl butyrate (S+) (Sigma-Aldrich, USA) and 0.1% of 40% heptanal+60% ethyl butyrate (S-) for 3-4 sessions for a total of 400-500 trials. The animals then rested for one to two weeks and were subsequently trained to discriminate the same odorant mixtures in sessions that took place 40 minutes after IP injection of 3mg/kg clozapine-N-oxide (CNO, Tocris, USA). 2) For the experiment in Supplementary Fig. 13 the animals were naïve for the discrimination of Iso vs. MO. The animals were injected intraperitoneally (IP) with 3mg/kg clozapine-N-oxide (CNO, Tocris, USA) 40 minutes before starting the session and were trained to discriminate between 1% Iso (S+) and MO (S-) for 6 sessions for a total of 500-600 trials. The reinforcement was then reversed and the animals were again injected with CNO 40 minutes before the sessions and were trained to discriminate between MO (S+) and 1% Iso (S-) for 5 sessions for a total of 400-500 trials.

### Two photon imaging of MLI activity in animals undergoing the Go-No Go task

All the animals were first habituated to the setup to minimize stress during the imaging experiments. All the imaging sessions started at least 10 minutes after mice had been head-fixed. We searched for active MLIs while imaging zones in the vermis of lobule VI, between the midline and the paravermal vein, an area of the cerebellum where GCs acquire a predictive feedback signal or expectation reward^29,30^. The head fixed two photon imaging system consisted of a movable objective microscope (MOM, Sutter Instrument Company, USA) paired with a 80 MHz, ∼100 femtosecond laser (Mai-Tai HP DeepSee, Spectra Physics, USA) centered at 920 nm. The MOM was fitted with a single photon epifluorescence eGFP filter path (475 nm excitation/500-550 nm emission) used for initial field targeting followed by switching to the two photon laser scanning path for imaging GCaMP at the depth of the MLIs. The galvometric laser scanning system was driven by SlideBook 6.0 (Intelligent Imaging Innovations, USA). The two photon time lapses were acquired at 256 x 256 pixels using a 1.0 NA/20x water emersion objective (Zeiss, Germany) at 5.3 Hz. On the day of initial imaging, a FOV was selected to image a large number of active cerebellar neurons located in the most superficial planes of the molecular layer (within 50 μm beyond the dorsal surface of the molecular layer) including mostly SCs, and several batches of 6000 frames (“time series”) were collected in each training session. After two photon imaging a second image of the vasculature was captured with wide field epifluorescence to reconfirm the field.

### Data analysis

Raw imaging data was first surveyed in ImageJ (NIH, USA) to exclude image sequences exhibiting axial movement. We did not find evidence of axial movement while the animal was engaged in the go-no go task. In addition, we performed control imaging where we excited GCaMP6f at 820 nm, a two photon excitation wavelength where fluorescence emission is Ca^2+^-independent^68^. Supplementary Fig. 16b,c shows that we did not detect transient changes in GCaMP6 fluorescence in a mouse engaged in the go-no go task when the cells were excited at 820 nm. If we found horizontal drift due to motion, we applied cross correlation-based image alignment using the Turboreg, Image J plugin. The data were then analyzed with CaImAn Matlab software that uses constrained nonnegative matrix factorization to define independent spatial and temporal components corresponding to changes in GCaMP fluorescence in individual MLIs^31^. Baseline of intensity (*F*0) was defined as the mean fluorescence intensity before trial start, defined as the time when the animal first licked. This was when fluorescence started to increase above baseline and the odorant was added at a random time 1-1.5 seconds after trial start. Intensity traces (*F*) were normalized according to the formula ΔF/F =(*F* − *F*0)/*F*0. After CaImAn analysis, the ΔF/F traces of the spatial components were sorted and we assigned trial traces to different behavioral events (Hit, CR, Miss and CR) and aligned them to trial start, odorant onset or water delivery. Finally, the time course for the average ΔF/F did not differ greatly between the different GCaMP variants as would be expected for a fast firing interneuron with small increases in Ca^2+^ per action potential (Supplementary Fig. 16a).

### Statistical analysis

Statistical analysis was performed in Matlab (Mathworks, USA). Statistical significance for changes in measured parameters for factors such as learning and odorant identity (S+ vs. S-) was estimated using a generalized linear model (GLM), with post-hoc tests for all data pairs corrected for multiple comparisons using false discovery rate^33^. The post hoc comparisons between pairs of data were performed either with a t-test, or a ranksum test, depending on the result of an Anderson-Darling test of normality. 95% CIs shown in the figures as vertical black lines or shading bounding the lines were estimated by bootstrap analysis of the mean by sampling with replacement 1000 times using the bootci function in MATLAB.

Principal component analysis was calculated using the Matlab Statistics Toolbox. Classification of trials using ΔF/F measured from all components in the FOV was accomplished via linear discriminant analysis (LDA) in Matlab. We obtained similar result with perceptron analysis (not shown). ΔF/F for all components for every trial except one were used to train the LDA, and the missing trial was classified by its fit into the pre-existing dataset. This was repeated for all trials and was performed separately for analysis where the identity of the odorants was shuffled. Fluorescence intensity traces, lick rates and kinematics in Fig. 6 were low-pass filtered with a hamming window of a time constant of 0.59 s.

### Dimensionality

Following Litwin-Kumar et al.^69^ we defined the dimension of the system (dim) with *M* inputs as the square of the sum of the eigenvalues of the covariance matrix of the measured ΔF/F for all ROIs in the FOV divided by the sum of each eigenvalue squared:

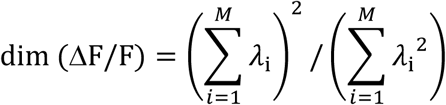

where λ_i_ are the eigenvalues of the covariance matrix of ΔF/F computed over the distribution of ΔF/F signals measured in the FOV. If the components of ΔF/F are independent and have the same variance, all the eigenvalues are equal and dim(ΔF/F) = *M*. Conversely, if the ΔF/F components are correlated so that the data points are distributed equally in each dimension of an *m*-dimensional subspace of the full *M*-dimensional space, only *m* eigenvalues will be nonzero and dim(ΔF/F) = *m*.

### Generalized linear modelling estimate of contribution of different variables to changes in ΔF/F

In order to quantify the contribution of different variables to neural activity, we used GLM as described by Engelhard et al. ^37^. We used the Matlab fitglm function to fit the per trial ΔF/F time course for mice proficient in the go-no go task with a GLM. We included “event” variables, “whole trial” variables and “continuous” variables. Continuous variables quantified kinematics including lick rate, the derivative of lick rate and the velocity and acceleration of movements made by base of the tail of the head fixed animal during the trial (body velocity and body acceleration, see Supplementary Fig. 3). The identity of the odorant (S+/S-odorant) was an event variable that increased from zero to one during the time for S+ or S-odorant application. Finally, whole trial variables were accuracy (1 for correct responses and 0 for incorrect responses), reinforcement history (1 for reinforcement in the last trial, 0 otherwise) and percent correct behavior calculated in a window of 20 trials.

### Data availability

The data supporting the findings of this study are available in GigaDB (http://gigadb.org/) (we are depositing the data in GigaDB, and this line will be replaced with a link to the data), the model is deposited in ModelDB (https://senselab.med.yale.edu/modeldb/)(we are depositing the data in ModelDB, and this line will be replaced with a link to the data) and the code is available in https://github.com/restrepd/CaImAnDR.

**Table 1.** Percent of ROIs displaying a change in ΔF/F during odorant application

| Calculation | Percent responsive | Number | Number of ROIs |
| --- | --- | --- | --- |
| Percent of ROIs responding in Hit trials | 99.9±0.23 | 21 time series, 5 mice | 74-199 |
| Percent of ROIs responding in CR trials | 82.3±26 | 21 time series, 5 mice | 74-199 |
| Percent of ROIs responding in Miss trials | 97.8±6.5 | 13 time series, 5 mice | 105-199 |
| Percent of ROIs responding in FA trials | 78±29 | 15 time series, 5 mice | 74-199 |
| Percent of ROIs responding in Miss trials that also respond in Hit trials | 99.96±0.15 | 13 time series, 5 mice | 105-199 |
| Percent of ROIs responding in FA trials that also respond in Hit trials | 99.8±0.47 | 15 time series, 5 mice | 74-199 |

This calculation was performed in time series that included at least one Miss or FA trial when the animal was proficient (percent correct >=80%). The ROI was classified as responsive when ΔF/F increased above or decreased below baseline ΔF/F by 2.5 x SD. Baseline mean and SD were calculated for the time interval 10 to 2 sec before odorant onset.

## References

1 Cayco-Gajic, N. A. & Silver, R. A. Re-evaluating Circuit Mechanisms Underlying Pattern Separation. Neuron 101, 584–602, doi:10.1016/j.neuron.2019.01.044 (2019).

2 Strick, P. L., Dum, R. P. & Fiez, J. A. Cerebellum and nonmotor function. Annu Rev Neurosci 32, 413–434, doi:10.1146/annurev.neuro.31.060407.125606 (2009).

3 Sokolov, A. A., Miall, R. C. & Ivry, R. B. The Cerebellum: Adaptive Prediction for Movement and Cognition. Trends Cogn Sci 21, 313–332, doi:10.1016/j.tics.2017.02.005 (2017).

4 Adamaszek, M. et al. Consensus Paper: Cerebellum and Emotion. Cerebellum 16, 552–576, doi:10.1007/s12311-016-0815-8 (2017).

5 Badura, A. et al. Normal cognitive and social development require posterior cerebellar activity. Elife 7, doi:10.7554/eLife.36401 (2018).

6 Jelitai, M., Puggioni, P., Ishikawa, T., Rinaldi, A. & Duguid, I. Dendritic excitation-inhibition balance shapes cerebellar output during motor behaviour. Nat Commun 7, 13722, doi:10.1038/ncomms13722 (2016).

7 Gao, Z., van Beugen, B. J. & De Zeeuw, C. I. Distributed synergistic plasticity and cerebellar learning. Nat Rev Neurosci 13, 619–635, doi:10.1038/nrn3312 (2012).

8 Marr, D. A theory of cerebellar cortex. J Physiol 202, 437–470 (1969).

9 Ito, M. Neural design of the cerebellar motor control system. Brain Res 40, 81–84, doi:10.1016/0006-8993(72)90110-2 (1972).

10 Albus, J. S. A theory of cerebellar function. Mathematical Biosciences 10, 25–61 (1971).

11 Heffley, W. & Hull, C. Classical conditioning drives learned reward prediction signals in climbing fibers across the lateral cerebellum. Elife 8, doi:10.7554/eLife.46764 (2019).

12 Tsutsumi, S. et al. Modular organization of cerebellar climbing fiber inputs during goal-directed behavior. Elife 8, doi:10.7554/eLife.47021 (2019).

13 Deverett, B., Koay, S. A., Oostland, M. & Wang, S. S. Cerebellar involvement in an evidence-accumulation decision-making task. Elife 7, doi:10.7554/eLife.36781 (2018).

14 Carta, I., Chen, C. H., Schott, A. L., Dorizan, S. & Khodakhah, K. Cerebellar modulation of the reward circuitry and social behavior. Science 363, doi:10.1126/science.aav0581 (2019).

15 Schonewille, M. et al. Reevaluating the role of LTD in cerebellar motor learning. Neuron 70, 43–50, doi:10.1016/j.neuron.2011.02.044 (2011).

16 Dean, P., Porrill, J., Ekerot, C. F. & Jorntell, H. The cerebellar microcircuit as an adaptive filter: experimental and computational evidence. Nat Rev Neurosci 11, 30–43, doi:10.1038/nrn2756 (2010).

17 Rancillac, A. & Crepel, F. Synapses between parallel fibres and stellate cells express long-term changes in synaptic efficacy in rat cerebellum. J Physiol 554, 707–720, doi:10.1113/jphysiol.2003.055871 (2004).

18 Liu, S. Q. & Cull-Candy, S. G. Synaptic activity at calcium-permeable AMPA receptors induces a switch in receptor subtype. Nature 405, 454–458, doi:10.1038/35013064 (2000).

19 Jorntell, H. & Ekerot, C. F. Receptive field plasticity profoundly alters the cutaneous parallel fiber synaptic input to cerebellar interneurons in vivo. J Neurosci 23, 9620–9631 (2003).

20 Jorntell, H. & Ekerot, C. F. Reciprocal bidirectional plasticity of parallel fiber receptive fields in cerebellar Purkinje cells and their afferent interneurons. Neuron 34, 797–806, doi:10.1016/s0896-6273(02)00713-4 (2002).

21 Jorntell, H., Bengtsson, F., Schonewille, M. & De Zeeuw, C. I. Cerebellar molecular layer interneurons - computational properties and roles in learning. Trends Neurosci 33, 524–532, doi:10.1016/j.tins.2010.08.004 (2010).

22 Liu, Y. et al. A single fear-inducing stimulus induces a transcription-dependent switch in synaptic AMPAR phenotype. Nat Neurosci 13, 223–231, doi:10.1038/nn.2474 (2010).

23 Wulff, P. et al. Synaptic inhibition of Purkinje cells mediates consolidation of vestibulo-cerebellar motor learning. Nat Neurosci 12, 1042–1049, doi:10.1038/nn.2348 (2009).

24 Ten Brinke, M. M. et al. Evolving Models of Pavlovian Conditioning: Cerebellar Cortical Dynamics in Awake Behaving Mice. Cell Rep 13, 1977–1988, doi:10.1016/j.celrep.2015.10.057 (2015).

25 Rowan, M. J. M. et al. Graded Control of Climbing-Fiber-Mediated Plasticity and Learning by Inhibition in the Cerebellum. Neuron 99, 999–1015 e1016, doi:10.1016/j.neuron.2018.07.024 (2018).

26 Gire, D. H., Whitesell, J. D., Doucette, W. & Restrepo, D. Information for decision-making and stimulus identification is multiplexed in sensory cortex. Nat Neurosci 16, 991–993, doi:10.1038/nn.3432 (2013).

27 Doucette, W. & Restrepo, D. Profound context-dependent plasticity of mitral cell responses in olfactory bulb. PLoS Biol 6, e258, doi:08-PLBI-RA-1962 [pii] 10.1371/journal.pbio.0060258 (2008).

28 Helmchen, F. & Denk, W. Deep tissue two-photon microscopy. Nature methods 2, 932–940, doi:10.1038/nmeth818 (2005).

29 Giovannucci, A. et al. Cerebellar granule cells acquire a widespread predictive feedback signal during motor learning. Nat Neurosci 20, 727–734, doi:10.1038/nn.4531 (2017).

30 Wagner, M. J., Kim, T. H., Savall, J., Schnitzer, M. J. & Luo, L. Cerebellar granule cells encode the expectation of reward. Nature 544, 96–100, doi:10.1038/nature21726 (2017).

31 Giovannucci, A. et al. CaImAn an open source tool for scalable calcium imaging data analysis. Elife 8, doi:10.7554/eLife.38173 (2019).

32 Chu, C. P., Bing, Y. H., Liu, H. & Qiu, D. L. Roles of molecular layer interneurons in sensory information processing in mouse cerebellar cortex Crus II in vivo. PLoS One 7, e37031, doi:10.1371/journal.pone.0037031 (2012).

33 Curran-Everett, D. Multiple comparisons: philosophies and illustrations. Am.J.Physiol Regul.Integr.Comp Physiol 279, R1–R8 (2000).

34 Gaffield, M. A. & Christie, J. M. Movement Rate Is Encoded and Influenced by Widespread, Coherent Activity of Cerebellar Molecular Layer Interneurons. J Neurosci 37, 4751–4765, doi:10.1523/JNEUROSCI.0534-17.2017 (2017).

35 Astorga, G. et al. Concerted Interneuron Activity in the Cerebellar Molecular Layer During Rhythmic Oromotor Behaviors. J Neurosci 37, 11455–11468, doi:10.1523/JNEUROSCI.1091-17.2017 (2017).

36 Tye, K. M. Neural Circuit Motifs in Valence Processing. Neuron 100, 436–452, doi:10.1016/j.neuron.2018.10.001 (2018).

37 Engelhard, B. et al. Specialized coding of sensory, motor and cognitive variables in VTA dopamine neurons. Nature 570, 509–513, doi:10.1038/s41586-019-1261-9 (2019).

38 Armbruster, B. N., Li, X., Pausch, M. H., Herlitze, S. & Roth, B. L. Evolving the lock to fit the key to create a family of G protein-coupled receptors potently activated by an inert ligand. Proc Natl Acad Sci U S A 104, 5163–5168, doi:10.1073/pnas.0700293104 (2007).

39 Gomez, J. L. et al. Chemogenetics revealed: DREADD occupancy and activation via converted clozapine. Science 357, 503–507, doi:10.1126/science.aan2475 (2017).

40 Raymond, J. L. & Medina, J. F. Computational Principles of Supervised Learning in the Cerebellum. Annu Rev Neurosci 41, 233–253, doi:10.1146/annurev-neuro-080317-061948 (2018).

41 Davie, J. T., Clark, B. A. & Hausser, M. The origin of the complex spike in cerebellar Purkinje cells. J Neurosci 28, 7599–7609, doi:10.1523/JNEUROSCI.0559-08.2008 (2008).

42 Otsu, Y. et al. Activity-dependent gating of calcium spikes by A-type K+ channels controls climbing fiber signaling in Purkinje cell dendrites. Neuron 84, 137–151, doi:10.1016/j.neuron.2014.08.035 (2014).

43 Rancz, E. A. & Hausser, M. Dendritic calcium spikes are tunable triggers of cannabinoid release and short-term synaptic plasticity in cerebellar Purkinje neurons. J Neurosci 26, 5428–5437, doi:10.1523/JNEUROSCI.5284-05.2006 (2006).

44 Eccles, J. C., Llinas, R. & Sasaki, K. The excitatory synaptic action of climbing fibres on the Purkinje cells of the cerebellum. J Physiol 182, 268–296, doi:10.1113/jphysiol.1966.sp007824 (1966).

45 Finch, E. A., Tanaka, K. & Augustine, G. J. Calcium as a trigger for cerebellar longterm synaptic depression. Cerebellum 11, 706–717, doi:10.1007/s12311-011-0314-x (2012).

46 Linden, D. J. & Connor, J. A. Long-term synaptic depression. Annu Rev Neurosci 18, 319–357, doi:10.1146/annurev.ne.18.030195.001535 (1995).

47 Brown, A. M. et al. Molecular layer interneurons shape the spike activity of cerebellar Purkinje cells. Sci Rep 9, 1742, doi:10.1038/s41598-018-38264-1 (2019).

48 Gaffield, M. A., Rowan, M. J. M., Amat, S. B., Hirai, H. & Christie, J. M. Inhibition gates supralinear Ca(2+) signaling in Purkinje cell dendrites during practiced movements. Elife 7, doi:10.7554/eLife.36246 (2018).

49 Jorntell, H. & Ekerot, C. F. Receptive Field Remodeling Induced by Skin Stimulation in Cerebellar Neurons in vivo. Front Neural Circuits 5, 3, doi:10.3389/fncir.2011.00003 (2011).

50 Lennon, W., Yamazaki, T. & Hecht-Nielsen, R. A Model of In vitro Plasticity at the Parallel Fiber-Molecular Layer Interneuron Synapses. Front Comput Neurosci 9, 150, doi:10.3389/fncom.2015.00150 (2015).

51 Szapiro, G. & Barbour, B. Multiple climbing fibers signal to molecular layer interneurons exclusively via glutamate spillover. Nat Neurosci 10, 735–742, doi:10.1038/nn1907 (2007).

52 Arlt, C. & Hausser, M. Microcircuit Rules Governing Impact of Single Interneurons on Purkinje Cell Output In Vivo. Cell Rep 30, 3020–3035 e3023, doi:10.1016/j.celrep.2020.02.009 (2020).

53 Bing, Y. H., Wu, M. C., Chu, C. P. & Qiu, D. L. Facial stimulation induces long-term depression at cerebellar molecular layer interneuron-Purkinje cell synapses in vivo in mice. Front Cell Neurosci 9, 214, doi:10.3389/fncel.2015.00214 (2015).

54 Chabrol, F. P., Blot, A. & Mrsic-Flogel, T. D. Cerebellar Contribution to Preparatory Activity in Motor Neocortex. Neuron 103, 506–519 e504, doi:10.1016/j.neuron.2019.05.022 (2019).

55 Hernandez-Briones, Z. S. et al. Olfactory stimulation induces cerebellar vermis activation during sexual learning in male rats. Neurobiol Learn Mem 146, 31–36, doi:10.1016/j.nlm.2017.11.003 (2017).

56 Sobel, N. et al. Odorant-induced and sniff-induced activation in the cerebellum of the human. J Neurosci 18, 8990–9001 (1998).

57 Mainland, J. D., Johnson, B. N., Khan, R., Ivry, R. B. & Sobel, N. Olfactory impairments in patients with unilateral cerebellar lesions are selective to inputs from the contralesional nostril. J Neurosci 25, 6362–6371, doi:10.1523/JNEUROSCI.0920-05.2005 (2005).

58 Kostadinov, D., Beau, M., Blanco-Pozo, M. & Hausser, M. Predictive and reactive reward signals conveyed by climbing fiber inputs to cerebellar Purkinje cells. Nat Neurosci 22, 950–962, doi:10.1038/s41593-019-0381-8 (2019).

59 Heffley, W. et al. Coordinated cerebellar climbing fiber activity signals learned sensorimotor predictions. Nat Neurosci 21, 1431–1441, doi:10.1038/s41593-018-0228-8 (2018).

60 Vincent, P. & Marty, A. Fluctuations of inhibitory postsynaptic currents in Purkinje cells from rat cerebellar slices. J Physiol 494 (Pt 1), 183–199, doi:10.1113/jphysiol.1996.sp021484 (1996).

61 Palay, S. L. & Chan-Palay, V. Cerebellar Cortex: Cytology and Organization. (Springer Berlin Heidelberg, 1973).

62 Dean, H. L., Hagan, M. A. & Pesaran, B. Only coherent spiking in posterior parietal cortex coordinates looking and reaching. Neuron 73, 829–841, doi:10.1016/j.neuron.2011.12.035 (2012).

63 Chen, T. W. et al. Ultrasensitive fluorescent proteins for imaging neuronal activity. Nature 499, 295–300, doi:10.1038/nature12354 (2013).

64 Astorga, G. et al. An excitatory GABA loop operating in vivo. Front Cell Neurosci 9, 275, doi:10.3389/fncel.2015.00275 (2015).

65 Kuhn, B., Ozden, I., Lampi, Y., Hasan, M. T. & Wang, S. S. An amplified promoter system for targeted expression of calcium indicator proteins in the cerebellar cortex. Front Neural Circuits 6, 49, doi:10.3389/fncir.2012.00049 (2012).

66 Li, A., Gire, D. H. & Restrepo, D. Y spike-field coherence in a population of olfactory bulb neurons differentiates between odors irrespective of associated outcome. J Neurosci 35, 5808–5822, doi:10.1523/JNEUROSCI.4003-14.2015 (2015).

67 Farneback, G. Two-frame motion estimation based on polynomial expansion. Proceedings of the 13th Scandinavian Conference on Image Analysis, 363–370 (2003).

68 Barnett, L. M., Hughes, T. E. & Drobizhev, M. Deciphering the molecular mechanism responsible for GCaMP6m’s Ca2+-dependent change in fluorescence. PLoS One 12, e0170934, doi:10.1371/journal.pone.0170934 (2017).

69 Litwin-Kumar, A., Harris, K. D., Axel, R., Sompolinsky, H. & Abbott, L. F. Optimal Degrees of Synaptic Connectivity. Neuron 93, 1153–1164 e1157, doi:10.1016/j.neuron.2017.01.030 (2017).

