## Supplementary Information for "Molecular layer interneurons in the cerebellum encode for valence in associative learning"

Ming Ma et al.

**Supplementary Note 1. The dimensionality of the MLI odorant responses is low.** The plots in Fig. 1c and Supplementary Fig. 4 indicate that odorant responses are alike between ROIs in the FOV. This raised the question whether the dimensionality of the MLI responses was small. In order to provide an estimate of the number of independent components comprising the ΔF/F responsiveness of the ensemble we calculated a quantitative measure of dimensionality[1] (see Methods). Supplementary Fig. 7 shows the dimensionality for four sessions with a total number of ROIs ranging from 103 to 136. The dimensionality ranged from 2 to 6 and did not differ significantly between time periods or between naïve and proficient (GLM p>0.05, 24 observations, 18 d.f., n=4 sessions, 4 mice, GLM F-statistic=2.42, p>0.05). This indicates that $Ca^{2+}$ ensemble activity is highly redundant for MLIs.

**Supplementary Note 2. LDA decoding analysis with subsets of ROIs.** Since dimensionality of MLI neural activity is low (Supplementary Fig. 7) a question arose as to whether performing LDA analysis on a fraction of the ROIs in the FOV would result in accurate decoding of the stimulus. We performed the LDA decoding analysis for smaller numbers of ROIs ranging from 1 to 100. We used pseudorandom sampling of fifty unique subsets of ROIs.  As expected, subsampling resulted in a decrease in accuracy of LDA decoding of the stimulus, but the decrease was relatively small and even a single ROI yielded a decoding accuracy significantly different from the shuffled control. In the experiment shown in Fig. 3e decoding accuracy decreases from 94% with all ROIs in the FOV to 83% with a single ROI, while the shuffled trials yield 50% accuracy. Fig. 3f shows the summary of this analysis for four sessions. A GLM analysis indicated that the decoding accuracy for the shuffled analysis was statistically different from accuracy with the subsets of ROIs ($p<0.001$, 72 observations, 54 d.f., n=4 sessions, 4 mice, GLM F-statistic=23.6, $p<0.001$). Thus, MLI activity encodes for the stimulus even when a single ROI is analyzed indicating that stimulus information encoded by MLI activity is highly redundant.

**Supplementary Note 3. MLI ensemble activity in error trials encodes the contextual identity of the odorant.** To gain a better understanding of the information on odorant valence present in the responses of the MLI ensemble we asked whether stimulus decoding accuracy calculated with LDA for proficient mice differed between correct (Hits and CRs), and incorrect trials (Miss and FAs). If information in MLI activity reflects the outcome of the trial we would expect that decoding accuracy would be lower for incorrect trials. On the other hand, if information encoded by MLI activity reflects the stimulus regardless of trial outcome decoding accuracy would not differ between correct and incorrect trials.

We performed this analysis for sessions that included at least one error trial (Miss or FA) when the animals were proficient. In these time series the majority of the ROIs exhibited changes in $\Delta F/F$ during the odorant application regardless whether the trial was a correct response (Hit or CR) or an error (Miss, FA) (Table 1). In addition, virtually all ROIs that responded with changes in $\Delta F/F$ during error trials also responded during Hit trials (Table 1, and see examples of $\Delta F/F$ time courses for single ROIs in Supplementary Fig. 9). As expected, in both the odorant application and reinforcement periods mean lick frequency for FA was higher than for CR and mean lick frequency for Miss trials was lower than Hits (Supplementary Fig. 8a, GLM analysis $p<0.001$, 48 observations, 40 d.f., n=6 sessions, 5 mice, GLM F-statistic=68, $p<0.001$, post-hoc t-test $p<pFDR=0.036$). In contrast, ΔF/F did not differ between Hits vs. Miss and CR vs. FA (Supplementary Fig. 8b and 8c). GLM analysis indicated that differences were not significant between Hits vs. Miss and CR vs. FA ($p>0.05$), while Hits/Miss differ from CR/FA ($p<0.01$, 48 observations, 40 d.f., n=6 sessions, 5 mice, GLM F-statistic=8.48, $p<0.001$, post-hoc t-test $p<pFDR=0.027$). To survey the information encoded in MLI activity in trials with different outcomes in proficient mice we utilized LDA analysis to decode the stimulus (Supplementary Fig. 8d: forward go-no go sessions, Supplementary Fig. 8e: reverse sessions). Decoding accuracy differed from shuffled for all outcomes and time periods (Supplementary Fig. 8d, t test, $p<pFDR=0.025$, n=6 sessions, 5 mice, Supplementary Fig. 8e t test, $p<pFDR=0.012$, n=3 sessions, 3 mice). In addition, GLM analysis did not find a significant difference between outcomes (Hit, Miss, CR or FA) or time period (odorant vs. reinforcement) indicating that MLI activity reflects the stimulus regardless of trial outcome (Supplementary Fig. 8d, $p>0.05$, 48 observations, 40 d.f., n=6 sessions, 5 mice, GLM F-statistic=1.49, $p>0.05$, Supplementary Fig. 8e, $p>0.05$, 24 observations, 16 d.f., n=3 sessions, 3 mice, GLM F-statistic=1.12, $p>0.05$). This analysis determined that odorant-induced MLI $Ca^{2+}$ changes carry information on the stimulus, as opposed to the outcome.

**Supplementary Note 4. Complementary analyses of the relationship between changes in ΔF/F and lick frequency.** We compared ΔF/F and lick frequency during the odorant application period for CR trials comparing the trials when the animal did not lick during the two 2 second odor response periods vs. CR trials when the animal licked (Supplementary Fig. 10). We did not find a difference between ΔF/F measured during the odorant period for the CR trials when the animal did not lick, compared to CR trials when the animal did lick during odorant application (Supplementary Fig. 10c, GLM yields no significant difference for the two types of CR trials, or for the different time periods, p>0.05, 36 observations, 30 d.f., n=6 sessions, 5 mice, GLM F-statistic=0.88, p>0.05).

Furthermore, we analyzed the relationship between the time course of ΔF/F and lick frequency in the time period shortly after odorant application when ΔF/F increases for both S+ and S-, before ΔF/F decreases for S- (and keeps increasing for S+). In Crus II, where neural activity of MLIs is thought to reflect licks, ΔF/F increases whenever there is an increase in licking frequency[2,3]. We aligned the traces to the point where the time derivative for ΔF/F increased above 0.03. We found that ΔF/F increased for both S+ and S- (Supplementary Fig. 11b, GLM analysis yields a significant change as a function of time, p<0.05, and no significant difference between S+ and S-, p>0.05, 132 observations, 128 d.f., 6 sessions, 5 mice, GLM F-statistic=3.1, p<0.05). In contrast, in this time period there was no increase in lick frequency (Supplementary Fig. 11a, GLM analysis yields no significant difference as a function of time, p>0.05 and a significant change between S+ and S-, p<0.001, 132 observations, 128 d.f., 6 sessions, 5 mice, GLM F-statistic=21, p<0.001). The data on the relationship between lick frequency and ΔF/F indicate that although there is a dependence between these two variables, the dependence is not consistent with a direct relationship between ΔF/F and lick frequency, as found in Crus II.

**Supplementary Note 5. Modeling the role of modulation of Purkinje cell output by molecular layer interneurons for the go-no go associative learning task.** In order to begin understanding the circuit basis of our findings in changes in lick behavior with chemogenetics in the go-no go associative learning olfactory discrimination task we generated a simple computational model of MLI interaction with PCs. Our model has a PC synaptically connected with two stellate cells (SCs) (Supplementary Fig. 14a). Both the SCs and the PC receive parallel fiber (PF) afferents. The odorant inputs increase the firing rate of PF inputs. The superficial SC inhibits the deep SC, and both SCs send inhibitory connections to the PC[4]. The PCs receive excitatory CF inputs in the dendrites and the SCs receive excitatory CF input through glutamate spillover. The CF inputs convey information about water reward. As a consequence, there is activation of the CF input after the presentation of the S+ odorant, but not during S- trials. Analogous to modeling of eye blink conditioning[5], we assumed that a pause in PC firing causes an increase in the lick rate that we modeled by convolution of the PC spikes with a reversed Gaussian function (see Supplementary Methods for details on the model). Simulation of S+ trials showed that odorant stimulus though PF inputs increases the activity of SCs that inhibit PC firing eliciting an increase in lick rate (Supplementary Fig. 14b, Saline S+). For the S- trials, we assumed the occurrence of a strong LTD at the PF-SC synapses and at the SC-PC synapses [6,7]. In this way, there is a reduction in the SC inhibition and an increase in PC excitation, yielding to a decrease in lick rate (Supplementary Fig. 14b, Saline S-).

To model the effect of inhibitory chemogenetics (Fig. 7 and Supplementary Fig. 13) we considered the effects of a reduction of 40% in the activity of the SC population. To model learning of the S+ odorant reward in the hM4Di+CNO condition, we reduced by 40% the synaptic weight of the synapses between PF-SC and SC-PC. This partial reduction still allowed the SCs to inhibit the PC and thus maintain a high lick rate (Supplementary Fig. 14b). Regarding the effect of learning on lick behavior for S- trials during the treatment hM4Di+CNO, we deemed that a 40% reduction in the activity of the SC population induced a diminished occurrence of LTD between the PF-SC and SC-PC synapses. The diminished LTD decreased the inhibition strength of SCs to pause the PC firing, which impaired the typical reduction of the lick rate during the S- condition (Supplemental. Fig. 14b). In summary, for the control condition we found an increase in lick strength for the S+ odorant that diverged from lick strength for S-. In contrast, for hM4Di+CNO lick strength increased slightly for both S+ and S-. A GLM analysis indicated that there were significant differences for lick strength for S+ vs S- ($p<0.01$) and CNO ($p<0.05$) and for the interactions between S+ vs. S- and CNO ($p<0.01$, 88 observations, 80 d.f., GLM F-statistic 7.9, $p<0.001$).

The model is simple. For example, it does not include basket cells. Furthermore, we did not model the large increase we find in the lick rate when the mouse receives the sugar water reward. Furthermore, we did not perform an exhaustive study of how the variables affect the changes in lick strength. Therefore, there are likely alternate explanations that will be explored in future studies with alternate computational models and an exhaustive search of the input variables. Regardless, our model provides a potential explanation for the results that will be tested in future experiments with slice electrophysiology and awake behaving recording.

### Supplementary Methods

Stellate cell simulation

We used reconstructed mouse SC morphology available in Neuromorpho (http://neuromorpho.org/neuron_info.jsp?neuron_name=GlyT2_030_Slice3_Stellate_cell). The morphology file was visualized using Blender with the addon NeuroMorphoVis[8]: https://github.com/BlueBrain/NeuroMorphoVis.

We removed the axons from the original swc morphology file and exported the reconstructed morphology into a NEURON hoc file (https://www.neuron.yale.edu/neuron/) using NLMorphologyViewer (http://www.neuronland.org). We proceeded to create the electrical compartmental model with passive and active properties of the SC membrane. The passive parameters of the SC model were adapted mainly from Molineux et al.[9]. We set the specific membrane resistivity $R_m = 20$ KΩ.cm$^2$, the specific membrane capacitance $C_m$=1.5 μF.cm$^{-2}$ [9], and the intracellular resistivity Ri=115 Ω.cm [10]. The input resistance Rin = 571.39 MΩ and membrane time constant $\tau_m$ = 40.30 ms were obtained injecting a hyperpolarizing current into the soma (-1 pA, 500ms). The time constant was obtained by a double exponential fit of membrane voltage decay. Those values are within the range of experimental values measured in SCs[9,11].

For modeling the active properties, we included voltage-dependent mechanisms for modeling the ionic channels at the soma of SCs. The firing patterns of SCs are regulated by fast sodium currents (Na), delayed rectifier potassium currents (KDR), A-type potassium currents (KA) and transient calcium currents (CaT)[9]. Since SCs and Golgi cells have similar firing properties[11], we adapted the voltage-dependent schemes of the conductances of Na, KDR, KA and CaT from a previous Golgi cell model[12] to reproduce the typical firing pattern of SCs[9] (Supplementary Fig. 15a).

Purkinje cell simulation

We adapted a previous two-compartment model that reproduces the typical spikes of PCs and is computationally efficient for constructing the cerebellum circuit model[13] (Supplementary Fig. 15b). The model was stimulated with background inhibitory inputs to present the typical curve of frequency versus current input from PCs[14].

Synaptic inputs

We used double exponential conductances to represent the synaptic inputs of the model with parameters taken from the literature.

PF – SC AMPARs[15]: $I_{max}$ = 96.42pA, $\bar{G}$ = 1.3774 nS, $\tau_1$ = 3.45 ms, $\tau_2$ = 3.17 ms, $E_{rev}$ = 0 mV.

PF – PC AMPARs[16]: $I_{max}$ = 20 pA, $\bar{G}$= 0.2857 nS, $\tau_1$ = 0.28 ms, $\tau_2$ = 1.23 ms, $E_{rev}$ = 0 mV.

SC-SC GABAaR[17,18]: $I_{max}$ = 75.5 pA, $\bar{G}$ = 1.0786 nS, $\tau_1$ = 0.6 ms, $\tau_2$ = 5.9 ms to 11.3 ms, $E_{rev}$ = -60 mV

SC-PC GABAaR[19]: $\bar{G}$ = 15 nS, $\tau_1$ = 1.8 ms, $\tau_2$ = 8.5 ms, $E_{rev}$ = -85 mV, Delay = 2 ms.

CF-PC. The CF inputs were modeled as a strong activation of the PC AMPARs.

CF-SC. We assume that glutamate spillover from CF to SC has a delay of ~10ms caused by glutamate diffusion[20] to stimulate the SC AMPARs. Based on the inferior olive neuron bursting patterns in response to CF inputs[21], we assumed a CF stimulus of 2 spikes at 300 Hz with an interval of 3.3 ms.

Synaptic plasticity

Synaptic plasticity was modeled by an increase or decrease of the maximum conductance of the synaptic channels. For modeling learning of S+ and S- tasks, we considered both the occurrence of LTD in SC-PC synapses[7], and the existence of LTD between the PF-SC synapses[6].

All simulations were performed on the NEURON simulator[22]. The model is deposited in ModelDB (https://senselab.med.yale.edu/modeldb/)(we are depositing the data in ModelDB, and this line will be replaced with a link to the data).

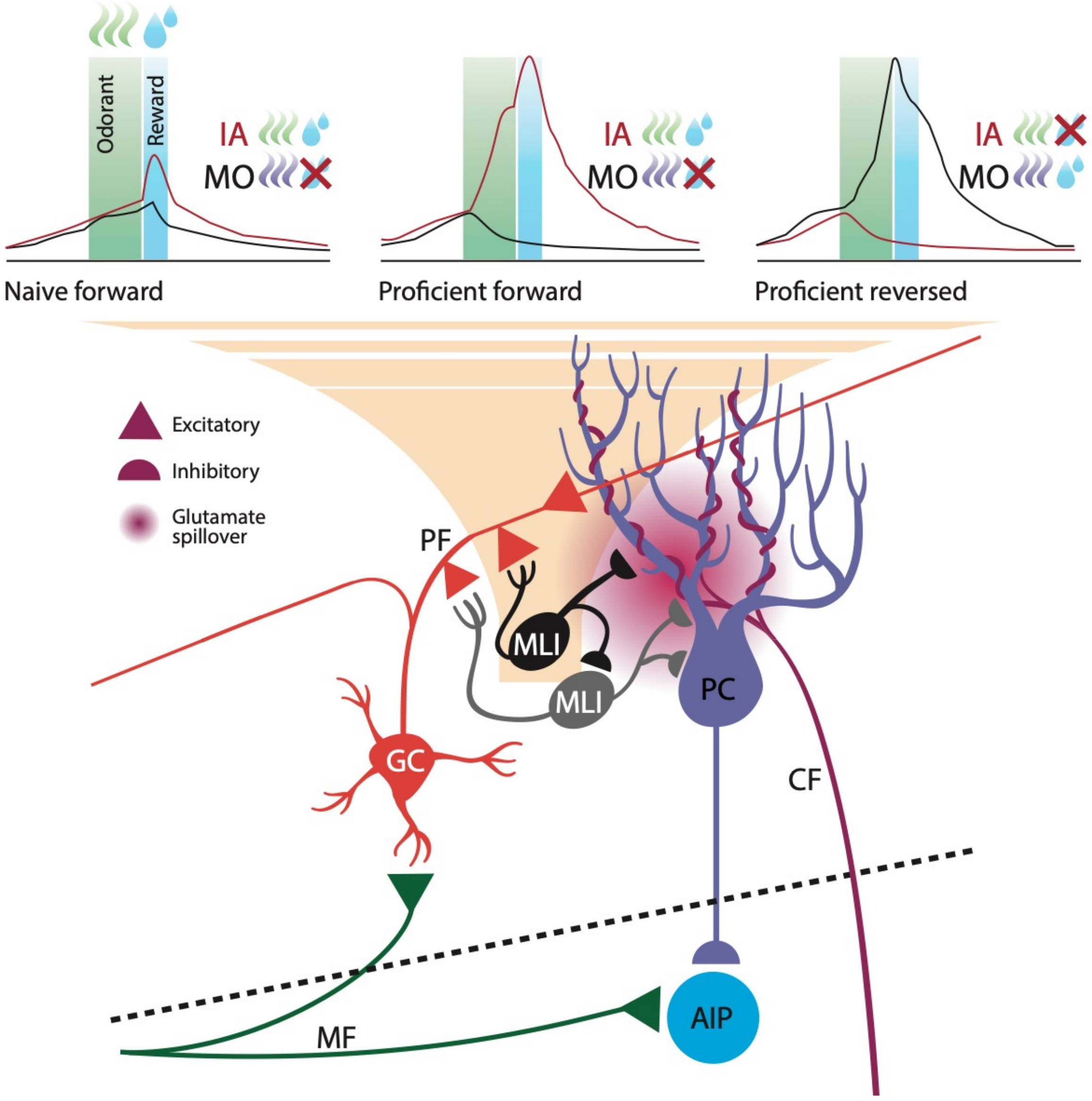


**Supplementary Figure 1. Diagram of the cerebellar circuit and summary of the findings.** This figure shows the circuit for the cerebellum. Climbing fibers (CFs) innervate the dendrites of Purkinje cells (PCs). Sensorimotor input is conveyed by Mossy fibers (MFs) to granule cells (GCs). The GCs innervate the dendrites of the PCs through parallel fibers (PFs). In this project we study the activity of molecular layer interneurons (MLIs) that receive innervation from GCs and inhibit PCs through a feedforward circuit. However, these neurons have also been proposed to from a disinhibitory motif where an MLI inhibits a deeper MLI that has a high probability of inhibiting PCs[4]. The traces on top represent the ΔF/F time courses recorded from MLIs in the naïve mouse (left), the proficient mouse (center) and the proficient mouse after reversal of the rewarded odorant (right). We find that the activity of the MLIs reflect the valence of the odorants.

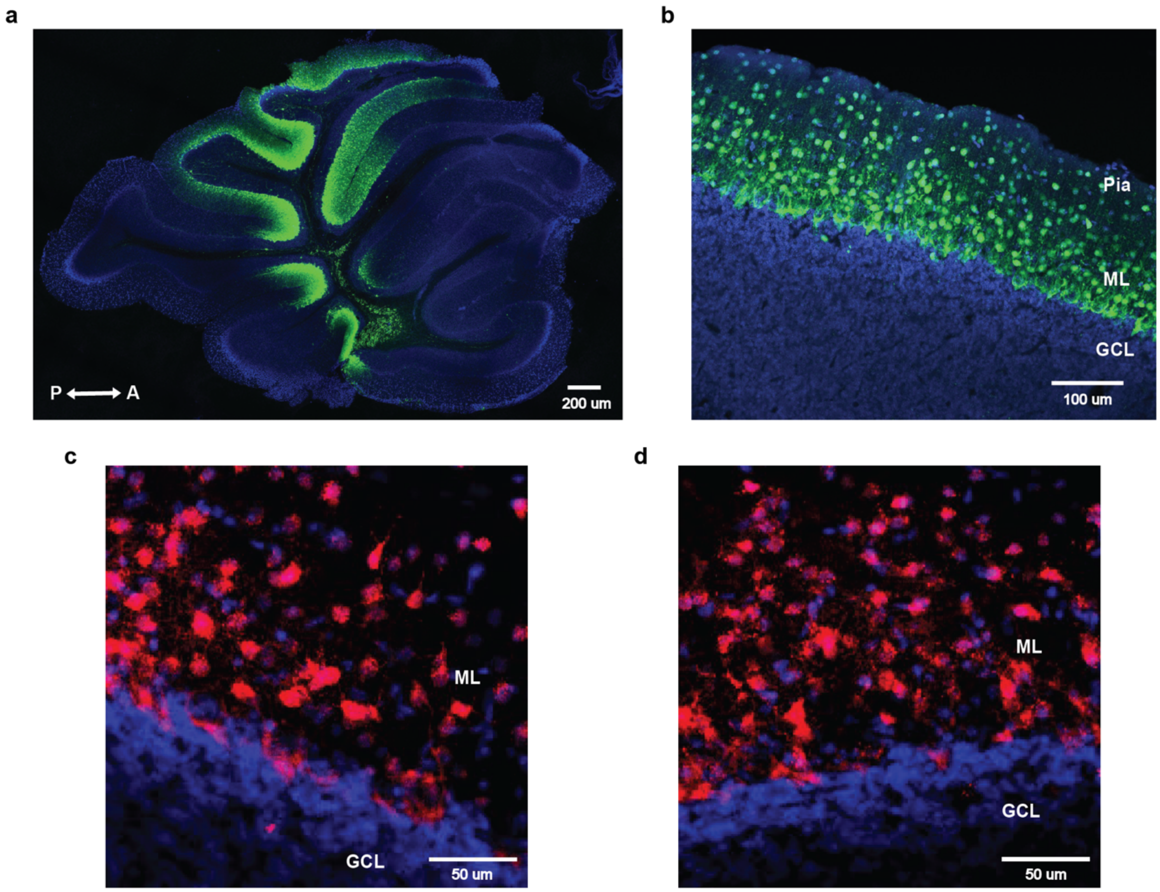


**Supplementary Figure 2. Expression of GCaMP6f and mCherry in fixed tissue slices. a,b.** GCaMP6f fluorescence in cerebellar sagittal brain slices from PV-Cre mice infected with AAV1-Syn-Flex-GCaMP6f. GCaMP6f (green) is expressed in the molecular layer. The slices were counterstained with DAPI.

**c,d.** mCherry fluorescence in cerebellar sagittal brain slices from PV-Cre mice infected with either AAV8-hSyn-DIO-mCherry (**c**) or AAV8-hSyn-DIO-hM4D(Gi)-mCherry (**d**). The slices were counterstained with DAPI.

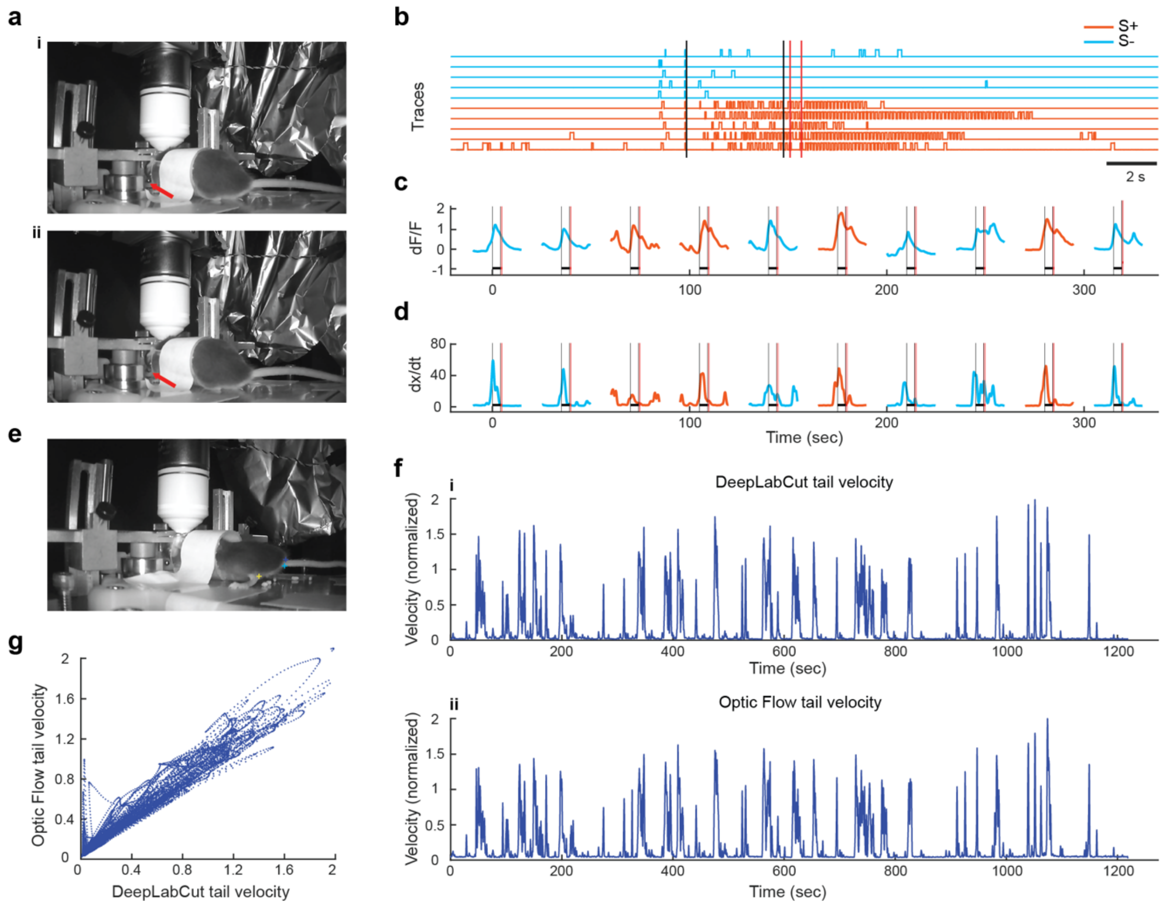


**Supplementary Figure 3. The mouse moves the body during the trial.**

**a.** Images show the mouse before (**i**) and at the start **(ii)** of the trial.

**b.** Lick traces for a subset of trials for a mouse that was proficient (>=80% correct) in the go-no go task. The mouse starts the trial by licking on the water spout and the odorant is delivered after a random delay of 1 to 1.5 sec. Orange: S+, light blue: S-. The two vertical lines denote times for odorant on and off (black lines) and reinforcement on and off (red lines).

**c.** Examples of ΔF/F $Ca^{2+}$ traces for a subset of trials.

**d.** Velocity of mouse movement for these trials measured using optical flow. The vertical black lines are odorant onset and removal and the red lines bound the reinforcement period.

**e.** DeepLabCut was used to track the base of the tail. Cross markers show the features tracked.

**f.** Comparison of the time course for the velocity of the tail measured with DeepLabCut (**i**) or optic flow (**ii**).

**g.** Relationship between the velocity of the tail measured with DeepLabCut vs. the velocity of the tail measured with optic flow for all time points in the traces shown in e and f. rho=0.98, $p<0.05$.

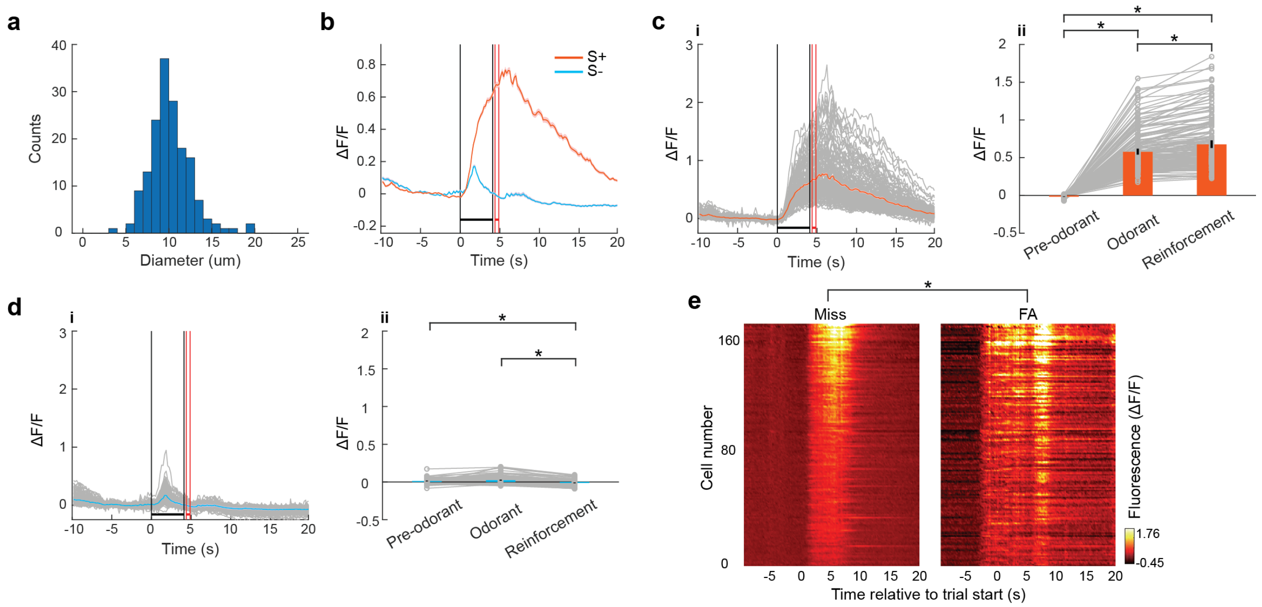


**Supplementary Fig. 4. Analysis of changes in ΔF/F for all regions of interest in the time stack for the example shown in Fig. 1.**

**a.** Histogram of the average diameter of the ROIs in Fig. 1biii. The average diameter was 10.5±4 µm (mean±SD, n=170).

**b.** Per trial time course for ΔF/F (mean±CI) for the S+ (orange) and S- (light blue) odorants. The vertical black lines are odorant onset and removal and the red lines bound the reinforcement period.

**c and d.** Per trial time course for ΔF/F for the S+ (ci) and S- (di) odorants. The orange and light blue lines are the mean ΔF/F calculated over all ROIs and the grey lines are per ROI time courses. cii and dii. ΔF/F calculated for 1 sec before the odorant (pre-odorant), the last second of odorant application (odorant) and 1.5 seconds after reinforcement (reinforcement) for the S+ (cii) and S- (dii) odorants. GLM analysis involving time periods and different odorants (S+ vs. S-) yielded significant differences for the interactions of reinforcement vs. pre-odorant and odorant vs. pre-odorant with S+ vs. S- ($p<0.001$, 1020 observations, 1014 degrees of freedom, n=170 ROIs, 1 mouse, GLM F-statistic 593, $p<0.001$). *Post-hoc ranksum p<pFDR=0.043.

**e.** Pseudocolor plots displaying the average per trial $\Delta F/F$ time course for FA and Miss for all ROIs in this example. GLM analysis involving time periods pre-odorant (1 sec before odorant onset), odorant (last second during odorant application) and reinforcement (1.5 seconds after reinforcement) and different events (Hits, Miss, CR and FA) yielded significant differences between reinforcement and pre-odorant ($p<0.001$), between odorant and pre-odorant ($p<0.001$), and between all interactions between these two period pairs and all events ($p<0.01$, 2040 observations, 2028 degrees of freedom, n=170 ROIs, 1 mouse, GLM F-statistic 234, $p<0.001$). *Post-hoc ranksum $p<pFDR=0.048$.

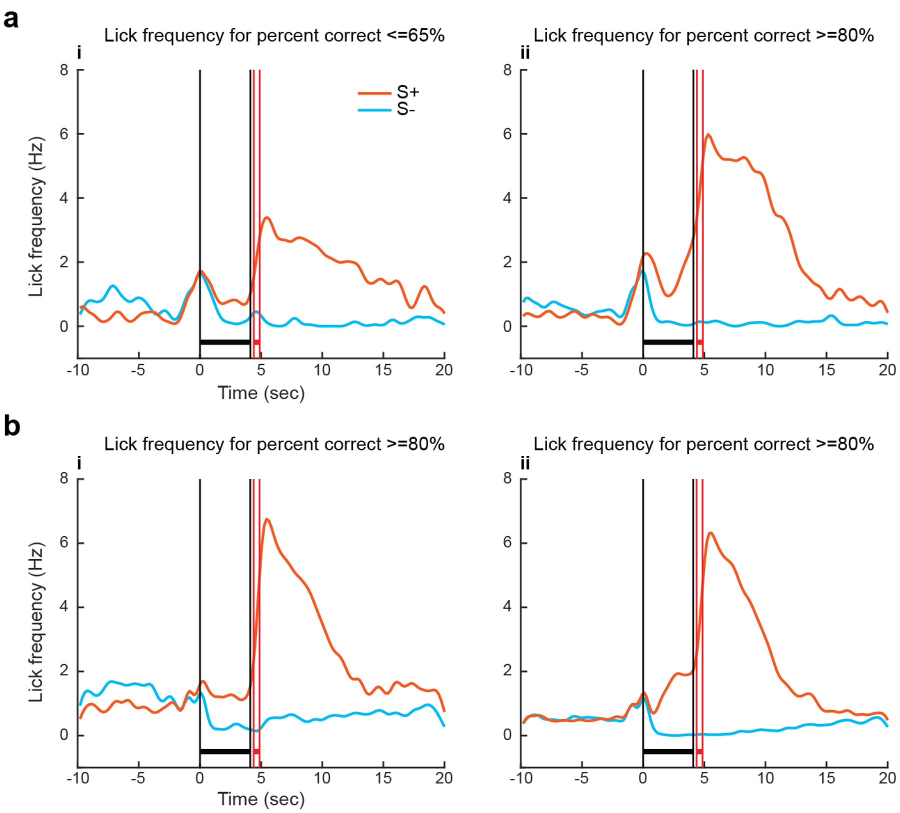


**Supplementary Figure 5. Mean lick frequency time courses shown for different experiments.**

**a.** Lick frequency time courses measured for the experiments whose decoding accuracy time courses are shown in Figs. 3ci and 3cii.

**b.** Lick frequency time courses measured for the experiments whose decoding accuracy time courses are shown in Figs. 4ei and 4eii. The vertical black lines are odorant onset and removal and the red lines bound the reinforcement period.

Orange: S+, light blue: S-.

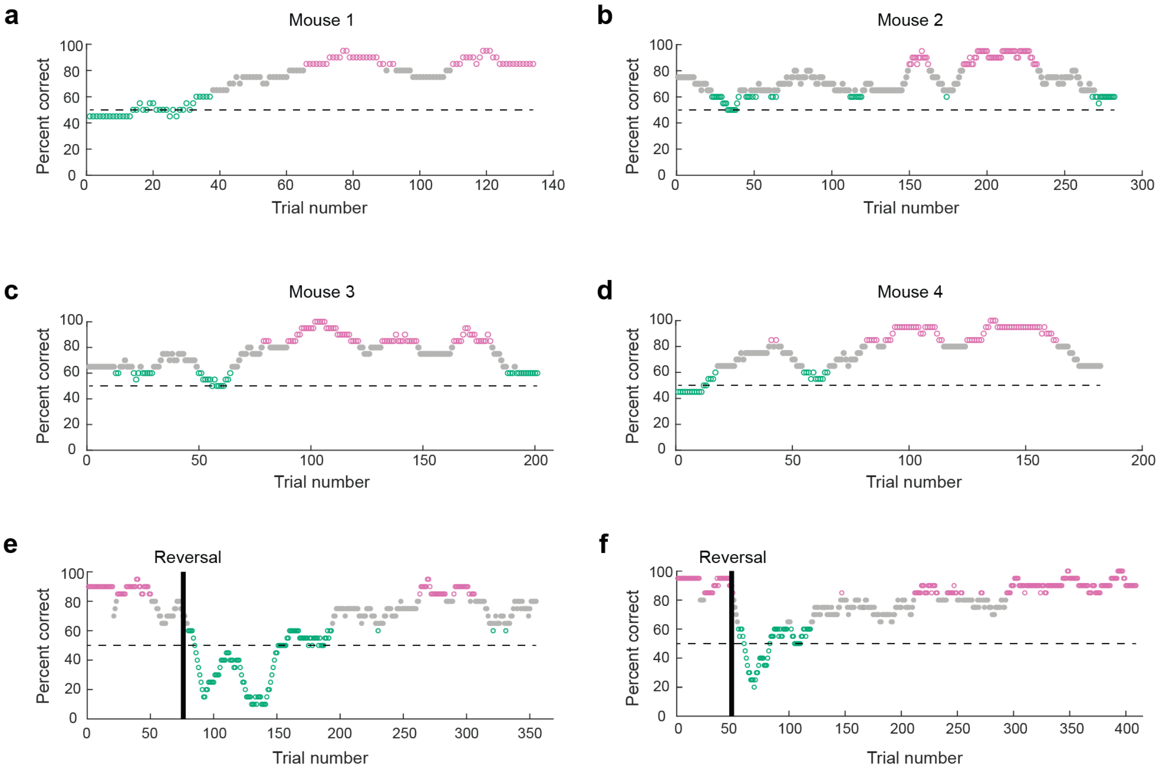


**Supplementary Figure 6. Behavioral performance for the mice included in the analysis in Figs. 2g and 4d,f.** The plots show percent correct performance for each mouse. Magenta: percent correct >=80%, green percent correct <=65%.

**a-d.** Percent correct for mice included in Fig. 2g.

**e-f.** Percent correct for two of the three mice included in Figs. 4d and f (the percent correct per trial for the other mouse is shown in Fig. 4a).

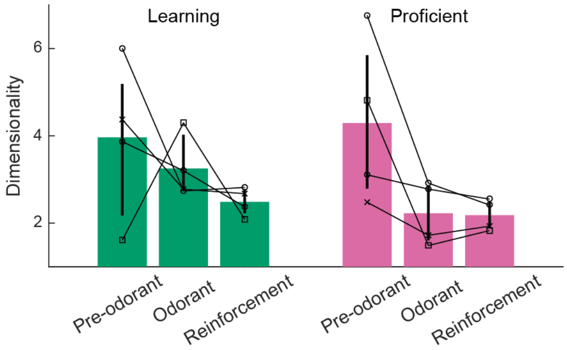


**Supplementary Figure 7. Dimensionality for ΔF/F.** Dimensionality was calculated in three periods: pre-odorant: 1 sec before odorant onset, odorant: last second of odorant application, reinforcement: 1.5 sec after the onset of water reward. Dimensionality was higher for the pre-odorant period compared to the odorant or reinforcement periods (GLM $p<0.01$, 18 d.f., n=4 sessions, 4 mice). Dimensionality did not differ between naïve and proficient, (GLM $p>0.05$, 24 observations, 18 d.f., n=4 sessions, 4 mice, GLM F-statistic=2.42, $p>0.05$).

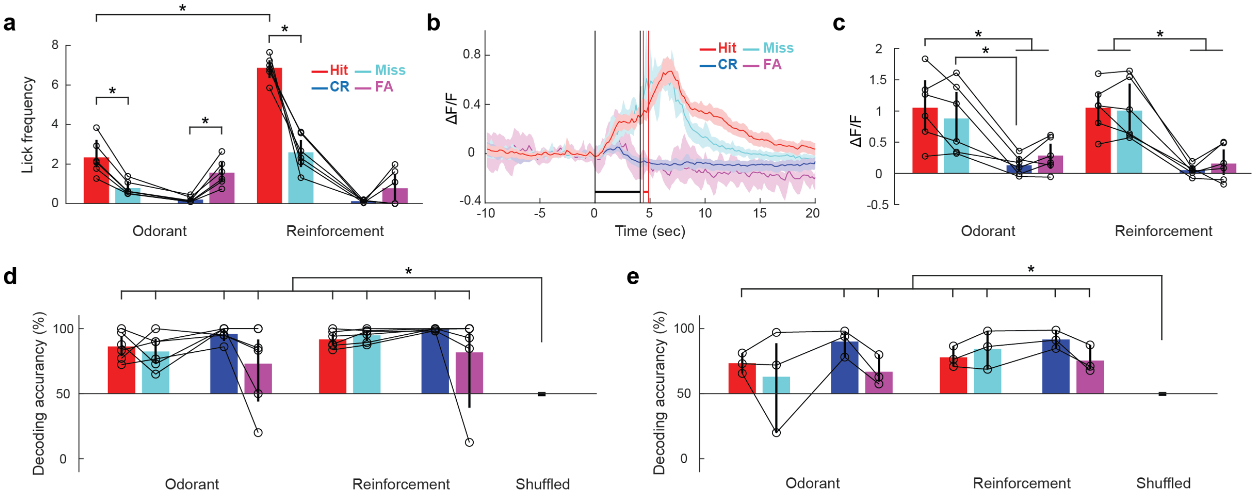


**Supplementary Figure 8. Linear discriminant analysis of stimulus identity for error trials for proficient mice.**

**a.** Mean lick frequency during the last second of odorant application and the first 1.5 seconds after reward delivery shown for the four trial outcomes. GLM analysis yields a statistically significant difference for odorant vs. reinforcement ($p<0.01$) and for the interaction between odorant vs. reinforcement and CR vs Hit ($p<0.01$) as well as for the interaction between odorant vs. reinforcement and FA vs Hit ($p<0.01$) (48 observations, 40 d.f., n=6 sessions, 5 mice, GLM F-statistic= 68, $p<0.001$). The color indicates the outcome of the trial (Red: Hit, Cyan: Miss, Blue: CR and Magenta: FA). *Post-hoc t-test $p<pFDR=0.036$.

**b**. Example of the time course for average ΔF/F (±CI, shade) for the four trial outcomes for one session for a mouse performing at percent correct >=80%. The vertical black lines are odorant onset and removal and the red vertical lines bound the reinforcement period. The corresponding lick rate time course is shown in Fig. 5b.

**c.** Mean ΔF/F during the last second of odorant application and the first 1.5 seconds after reward delivery shown for the four trial outcomes. GLM analysis yields a statistically significant difference for CR vs Hit ($p<0.001$) as well as for FA vs Hit ($p<0.01$) (48 observations, 40 d.f., n=6 sessions, 5 mice, GLM F-statistic=8.48, $p<0.001$). *Post-hoc t-test, $p<pFDR=0.027$. The lick time course corresponding to this ΔF/F time course is shown in Fig. 5b

**d and e.** Decoding accuracy for LDA analysis of stimulus prediction calculated with ΔF/F for all ROIs in the FOV classified by trial outcome. **d.** Forward sessions where S+ was Iso and S- was MO. Decoding accuracy differed from shuffled for all outcomes and time periods (t test, $p<pFDR=0.011$, n=6 sessions, 5 mice). GLM did not yield a significant difference for score categories or time period (odorant vs. reinforcement) (48 observations, 40 d.f. $p>0.05$, n=6 sessions, 5 mice, GLM F-statistic=1.49, $p>0.05$). A two-sample *F*-test for equal variances indicated that the variance differs between FA and CR ($p<pFDR=0.025$, n=6 sessions, 5 mice). *Post-hoc t-test, $p<pFDR=0.01$. **e.** Reverse sessions where S+ was MO and S- was Iso. Decoding accuracy differed from shuffled for all outcomes and time periods (t test, $p<pFDR=0.012$, n=3 sessions, 3 mice). GLM did not yield a significant difference for score categories or time period (odorant vs. reinforcement) (24 observations, 16 d.f. $p>0.05$, n=3 sessions, 3 mice, GLM F-statistic=1.12, $p>0.05$). *Post-hoc t-test, $p<pFDR=0.013$.

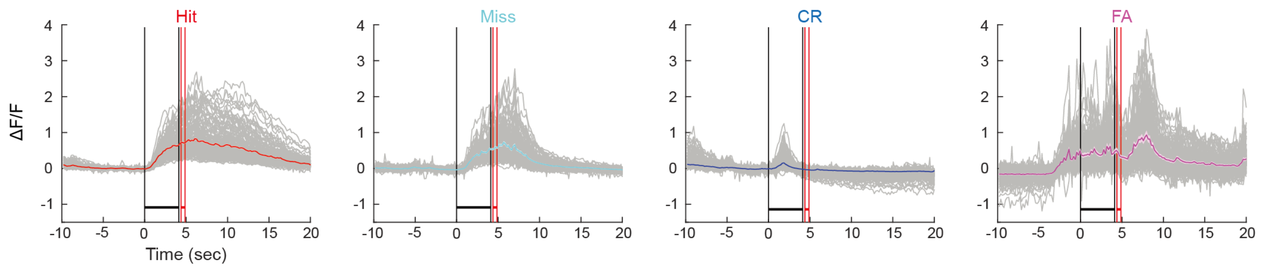


**Supplementary Figure 9. Examples of time courses of ΔF/F for different behavioral outcomes in proficient mice.**

Examples of ΔF/F time courses for 155 ROIs in the FOV shown. The ΔF/F time courses are shown as an average for all trials belonging to specific behavioral outcomes (Hit, Miss, CR or FA) for a single time series.

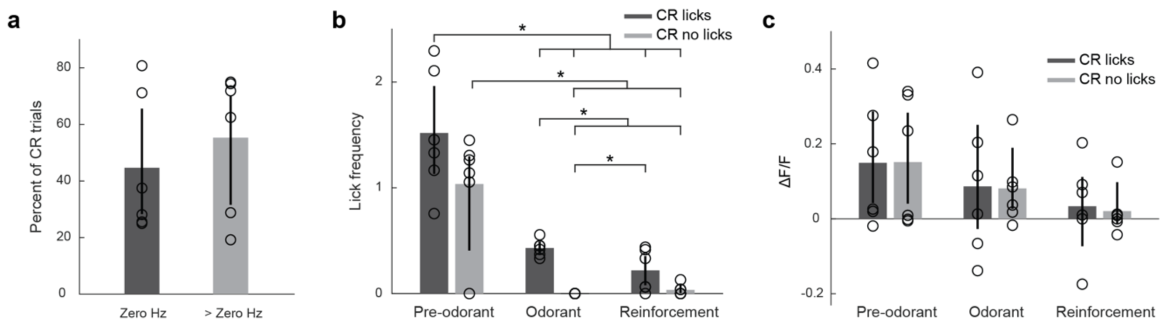


**Supplementary Figure 10. Lick frequency and ΔF/F for CR trials with zero licks for proficient mice.** The CR trials were classified between trials with no licks in the two 2 second segments for evaluation of lick responses during odorant application (CR-no licks), and other CR trials (CR-licks).

**a.** Percent of CR-no licks trials vs. CR-licks trials. A t test does not find a significant difference (p>0.05, n=6 sessions, 5 mice).

**b.** Lick frequency for CR-no licks and CR-licks for three different periods: one second before odorant application (Pre-odorant), during the two 2 second segments (Odorant) and for 1.5 seconds following reinforcement (Reinforcement). A GLM analysis found a statistically significant difference between CR-licks and CR-no licks (p<0.05), and odorant vs. pre-odorant and reinforcement vs. pre-odorant (p<0.001, 36 observations, 30 d.f., n=6 sessions, 5 mice, GLM F-statistic=20.3, p<0.001).

**c.** ΔF/F for the same time periods. A GLM found no differences between the two CR trials and between time periods (p>0.05, 36 observations, 30 d.f., n=6 sessions, 5 mice, GLM F-statistic 0.88, p>0.05).

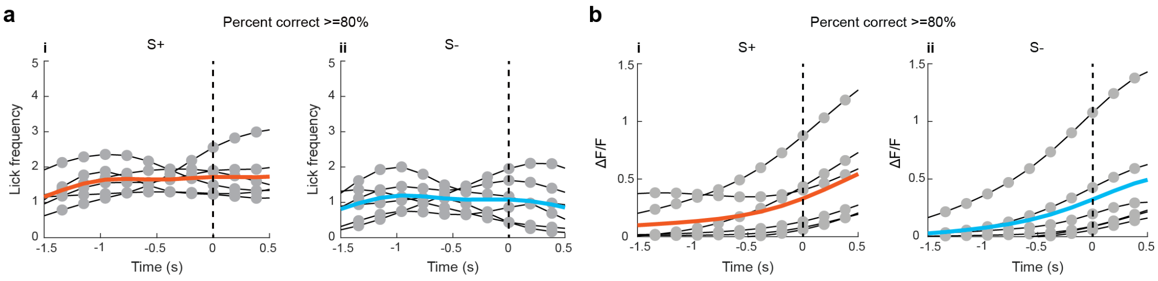


**Supplementary Figure 11. Mean ΔF/F and lick frequency time courses for the time period shortly after odorant application when ΔF/F starts increasing for both the S+ and S- odorants.** The time courses were aligned to the time after odorant addition when the derivative of ΔF/F increased above 0.03. The data are for six proficient mice. Mean values are shown as a thick orange or light blue line.

**a.** Lick frequency calculated as the mean over all trials when the mouse was proficient. In contrast, in this time period there was no increase in lick frequency. GLM analysis yields no significant difference as a function of time, $p>0.05$ and a significant change between S+ and S-, $p<0.001$, 132 observations, 128 d.f., n=6 sessions, 5 mice, GLM F-statistic=21, $p<0.001$. **i.** S+ odorant. **ii.** S- odorant.

**b.** ΔF/F calculated as the mean over all trials when the mouse was proficient. GLM analysis yields a significant change as a function of time, $p<0.05$, and no significant difference between S+ and S-, $p>0.05$, 132 observations, 128 d.f., n=6 sessions, 5 mice, GLM F-statistic = 3.1, $p<0.05$.

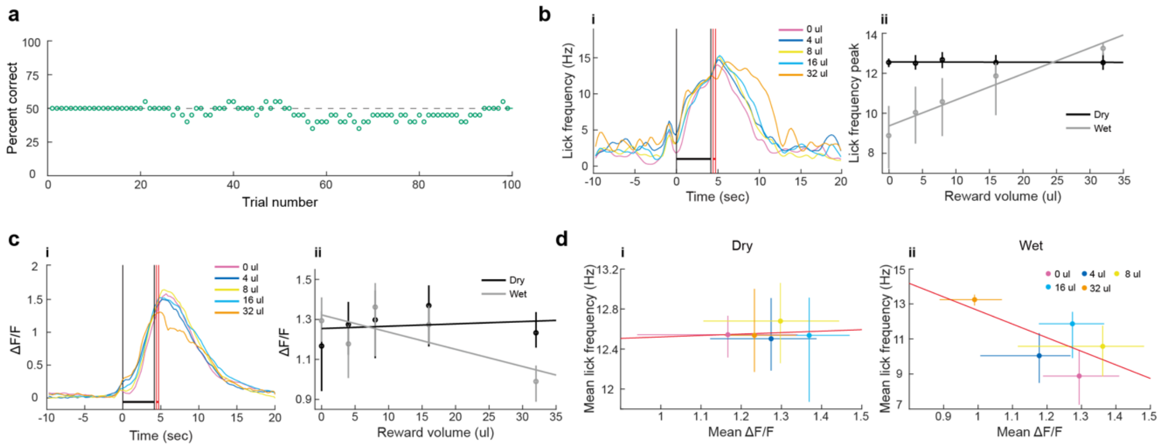


**Supplementary Fig. 12. Evaluation of changes in ΔF/F responses for MLIs when the volume of sugar water reward is varied.** Results are shown for ΔF/F responses of MLIs and lick frequency for one animal engaged in a go-go task where both odorants were rewarded equally. The volume of sugar water delivered for successful trials was varied from 0 to 32 µl.

**a.** Percent correct behavior scored as if this were a go-no go task shows that the mouse responded to both odorants in the go-go task (~50% correct response).

**b. i.** Time course for lick frequency in this go-go experiment. **ii.** Mean per trial lick frequency (±95% CI) calculated at two time points: 4 sec (when the animal is doing dry licks, dry time point) and at 7.5 sec (when the animal is licking to receive the reward, wet timepoint). GLM analysis yielded a statistically significant difference for lick frequency for dry vs. wet licking and for the interaction between the volume of sugar water delivered and dry vs. wet ($p<0.001$, 198 observations, 194 d.f., 1 session, 1 mouse, GLM F-statistic=21.8, $p<0.001$).

**c. i.** Time course for the mean ΔF/F for all trials (averaged for all ROIs per trial in the FOV). **ii.** Mean ΔF/F (±95% CI) calculated at the dry and wet time points. GLM analysis did not yield statistically significant differences ($p>0.05$, 198 observations, 194 d.f., 1 session, 1 mouse, GLM F-statistic=1.7, $p>0.05$).

**d.** Relationship between the mean lick frequency and mean ΔF/F for dry (**i**) and wet (**ii**) time points. The correlation coefficients were not significant for dry ($\rho$=0.16, $p>0.05$) or wet licking ($\rho$=-0.66, $p>0.05$).

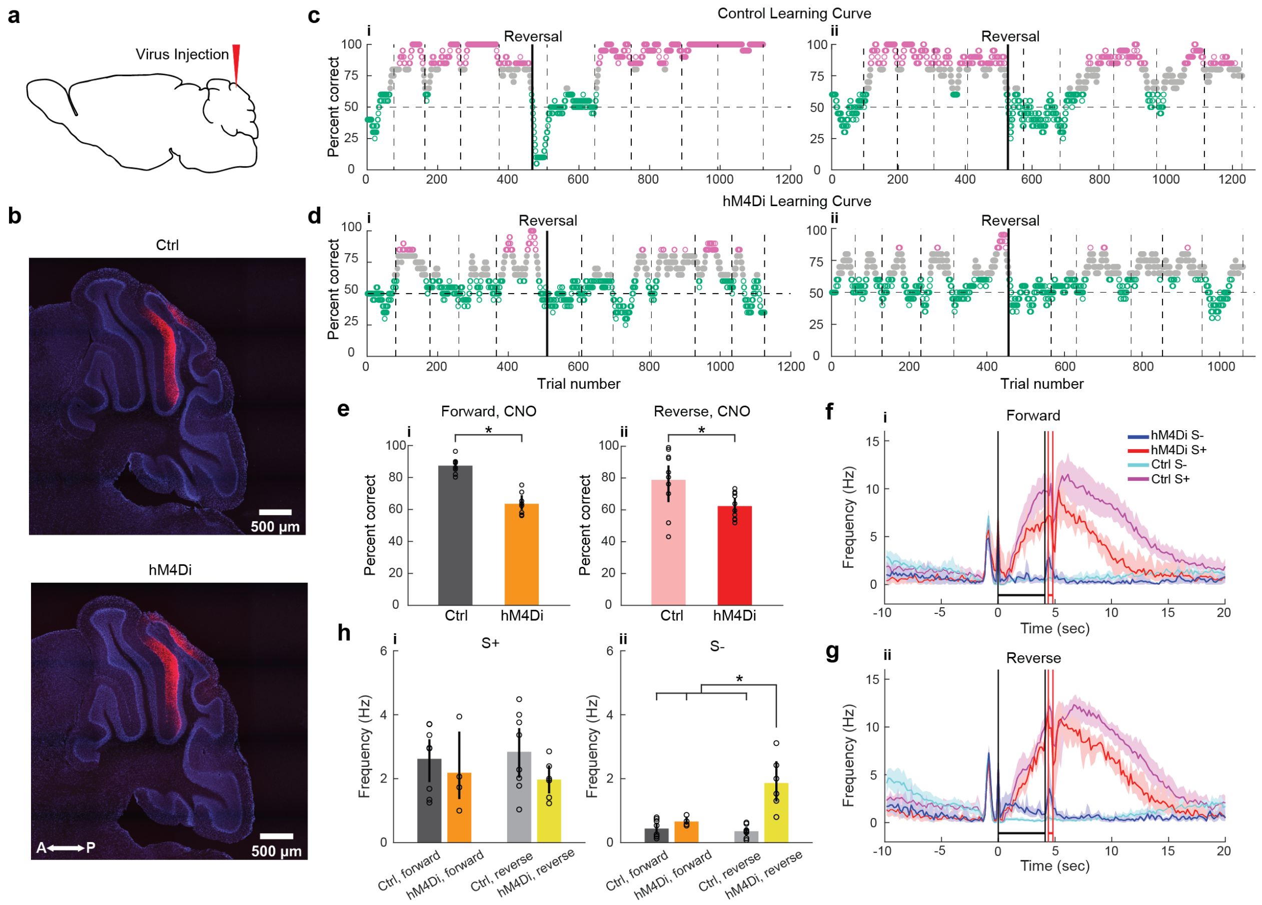


**Supplementary Fig. 13. Chemogenetic inhibition of MLI activity impairs associative learning.**

**a.** Location of virus injection shown in a sagittal diagram of the brain.

**b.** Expression of mCherry and DAPI in the cerebellum 2 months after injection of either AAV8-hSyn-DIO-hM4D(Gi)-mCherry (bottom) or AAV8-hSyn-DIO-mCherry (top) in PV-Cre mice.

**c. and d.** Examples of behavioral performance in a go-no go task for mice that learned to differentiate between Iso and MO. Green: <=65% correct, magenta: >=80% correct. For these experiments CNO was injected before the start of the session. At the beginning of the experiment (forward session) the rewarded odorant (S+) was 1% iso-amyl acetate and the unrewarded odorant was S-: 1% mineral oil. The reward was reversed at the trial denoted by the vertical line (reversed session). **c.** Control mice expressing mCherry in MLIs. **d.** Mice expressing hM4Di in MLIs. **i** and **ii** show the results of experiments with two separate mice.

**e.** Mean percent correct for behavioral performance (mean$\pm$ 95% CIs, n=4 sessions forward, 5 sessions reversed, 2 mice) for forward (**i**) and reversed (**ii**) sessions. Note: the first session was excluded because that is usually a low percent session when the animal is learning. A GLM analysis indicates that there is a difference between genotypes ($p<0.001$, 36 observations, 32 d.f., 4 sessions for hM4Di and 5 sessions for control, 2 mice, GLM F-statistic=9.3, $p<0.001$).

**f**. Average lick frequency time course for forward (**i**) and reversed (**ii**) sessions for mice performing >75% correct (mean$\pm$ 95% CI, shade). The vertical black lines are odorant onset and removal and the red lines bound the reinforcement period.

**g**. Mean lick frequency ($\pm$CI) during the initial portion of the odorant application period (0.8 to 1.8 sec) for mice performing >75% correct (**i**: S+, **ii**: S-). GLM analysis did not find a statistically significant difference for treatment or hM4Di expression (or interactions) for S+ ($p>0.05$, 26 observations, 22 d.f., two hM4Di mice and two control mice, 8 sessions, GLM F-statistic=0.92, $p>0.05$), and found a difference for genotype x forward vs. reversed for S- ($p<0.001$, 26 observations, 22 d.f, two hM4Di mice and two control mice, 8 sessions, GLM F-statistic=16.2, $p<0.001$). *$p<p_{FDR}=0.03$ for post-hoc t-test.

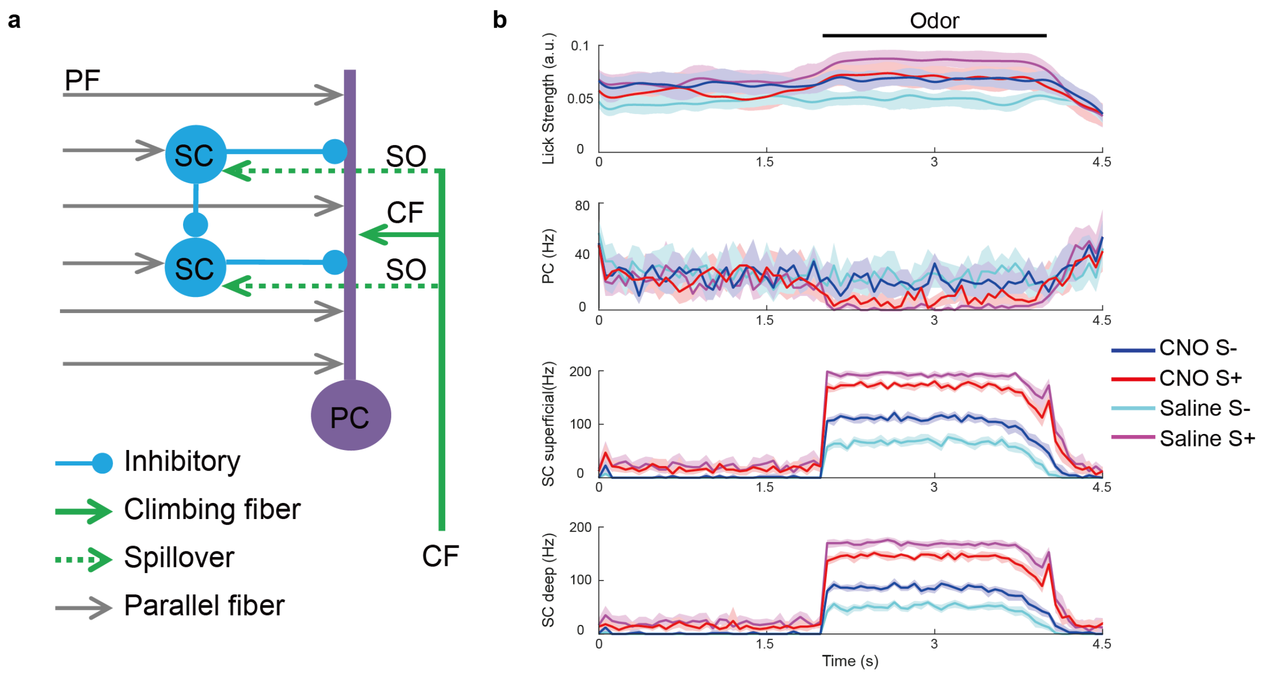


**Supplementary Figure 14. Computational simulation of the modulation of PC activity by MLIs. a.** Schematic representation of the model of the SC/PC circuit. Both SCs and the PC receive excitatory PF inputs. The SCs send inhibitory inputs to the PC. The superficial SC also sends inhibitory inputs to the deep SC. The PC receives strong excitatory inputs from CFs. The SCs also receive excitatory inputs through glutamate spillover from CFs. **b.** Results of odorant stimulation of the neural circuit. The odorant inputs are represented as a sustained increase in the PF firing rate from 2 to 4 seconds. From the top to the bottom, the panels represent lick strength, PC firing rate, and superficial and deep SC firing rates. The model's response for each experimental condition is represented by a specific color. A GLM analysis indicated that there were significant differences for lick strength for S+ vs S- ($p<0.01$) and CNO ($p<0.05$) and for the interactions between S+ vs. S- and CNO ($p<0.01$, 88 observations, 80 d.f., GLM F-statistic 7.9, $p<0.001$).

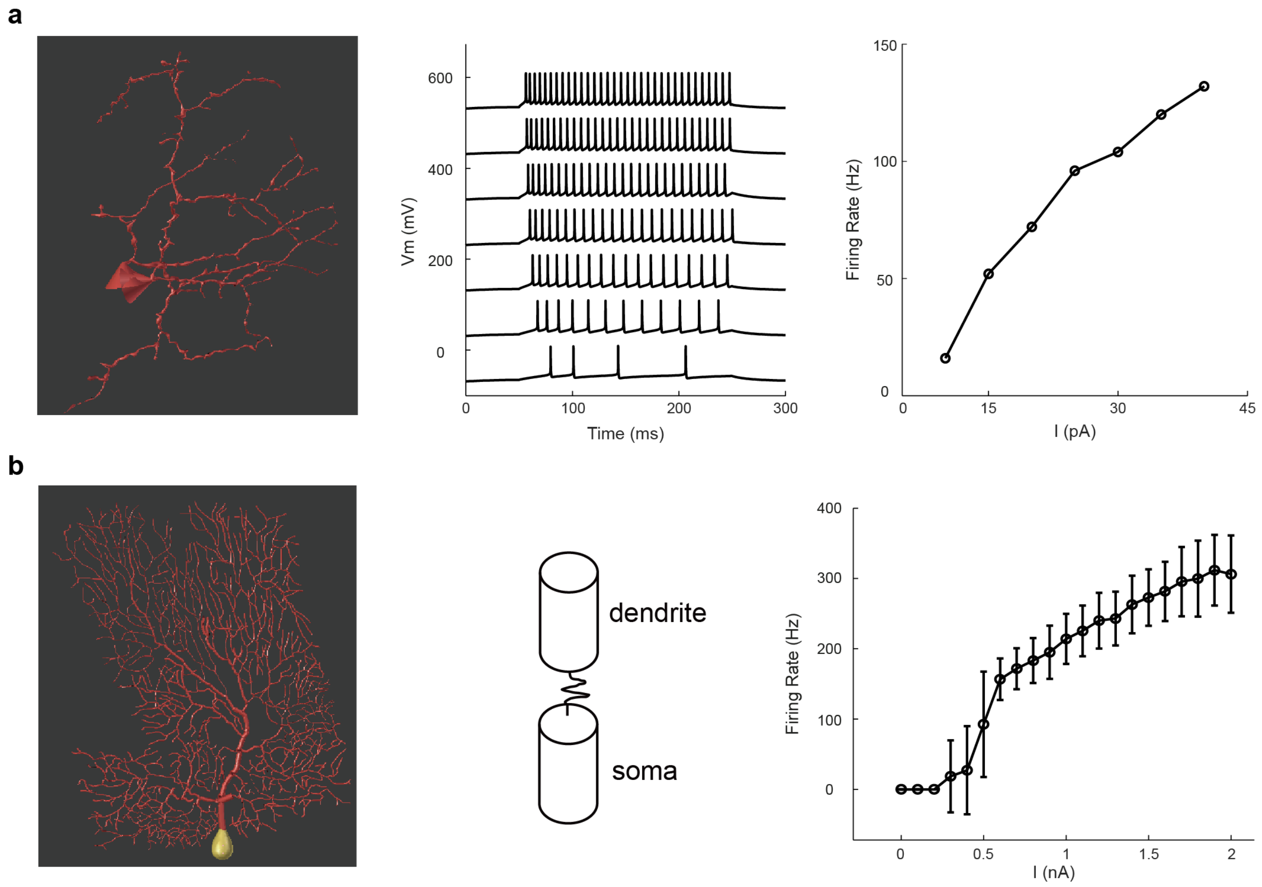


**Supplementary Figure 15. Compartmental models of SCs and PC. a.** Compartmental model of SC. The model presents the typical firing response of SCs to increasing intensities of a step current of 200ms. **b.** The reduced two compartmental model of PC[13] in the presence of background inhibitory inputs reproduces the typical curve of firing frequency versus input current[14] . The representative full morphology of the PC was taken from Martone and colleagues[23].

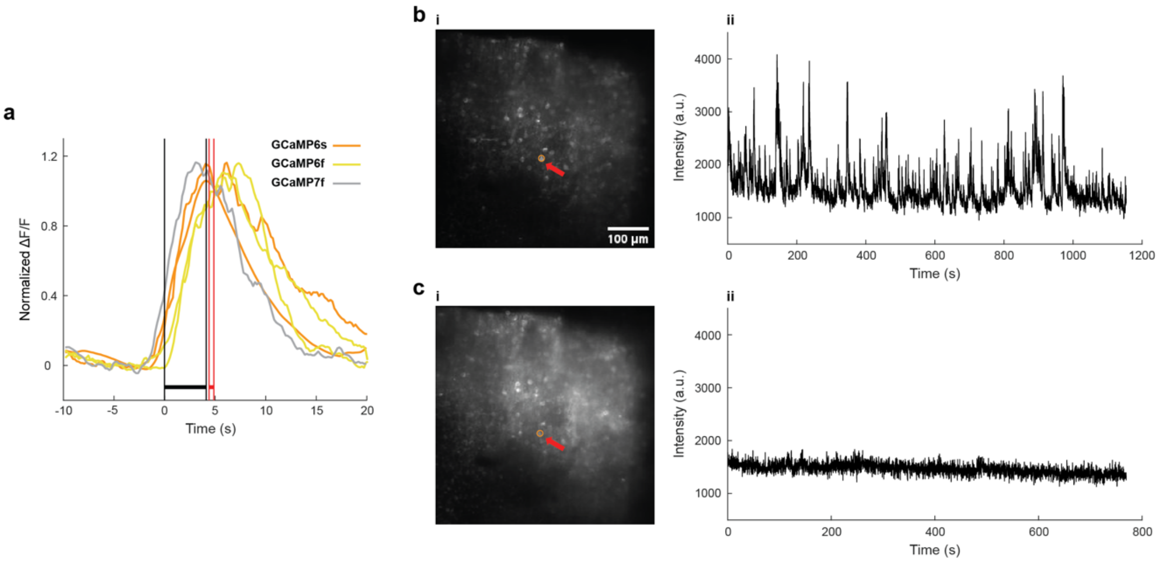


**Supplementary Figure 16. Controls for ΔF/F measurements of MLI cytoplasmic $Ca^{2+}$ with GCaMP in the animal engaged in the go-no go task.**

**a.** Time courses for average ΔF/F measured for rewarded odorant trials in MLIs of five proficient mice expressing different GCaMP proteins.

**b and c.** Control showing that the changes in ΔF/F are not due to axial movements of the ROIs. Two photon imaging was performed in the molecular layer of a mouse engaged in the go-no go task with excitation at either 920 nm (a, GCaMP6f emission $Ca^{2+}$-sensitive) or 820 nm (b, GCaMP6f emission $Ca^{2+}$-insensitive). Fluorescence transients were detected at 920 nm excitation, but not at 820 nm excitation indicating that the transients were not due to axial movement.

**Supplementary Movie 1.** The movie shows mouse movement for 29 seconds starting ~10 seconds before trial initiation.

**Supplementary Movie 2.** The movie shows changes in GCaMP6f fluorescence for the MLIs in Fig. 1b.

### Supplementary References


1 Litwin-Kumar, A., Harris, K. D., Axel, R., Sompolinsky, H. & Abbott, L. F. Optimal Degrees of Synaptic Connectivity. *Neuron* **93**, 1153-1164 e1157, doi:10.1016/j.neuron.2017.01.030 (2017).

2 Gaffield, M. A. & Christie, J. M. Movement Rate Is Encoded and Influenced by Widespread, Coherent Activity of Cerebellar Molecular Layer Interneurons. *J Neurosci* **37**, 4751-4765, doi:10.1523/JNEUROSCI.0534-17.2017 (2017).

3 Astorga, G. *et al.* Concerted Interneuron Activity in the Cerebellar Molecular Layer During Rhythmic Oromotor Behaviors. *J Neurosci* **37**, 11455-11468, doi:10.1523/JNEUROSCI.1091-17.2017 (2017).

4 Arlt, C. & Hausser, M. Microcircuit Rules Governing Impact of Single Interneurons on Purkinje Cell Output In Vivo. *Cell Rep* **30**, 3020-3035 e3023, doi:10.1016/j.celrep.2020.02.009 (2020).

5 ten Brinke, M. M. *et al.* Evolving Models of Pavlovian Conditioning: Cerebellar Cortical Dynamics in Awake Behaving Mice. *Cell Rep* **13**, 1977-1988, doi:10.1016/j.celrep.2015.10.057 (2015).

6 Lennon, W., Yamazaki, T. & Hecht-Nielsen, R. A Model of In vitro Plasticity at the Parallel Fiber-Molecular Layer Interneuron Synapses. *Front Comput Neurosci* **9**, 150, doi:10.3389/fncom.2015.00150 (2015).

7 Bing, Y. H., Wu, M. C., Chu, C. P. & Qiu, D. L. Facial stimulation induces long-term depression at cerebellar molecular layer interneuron-Purkinje cell synapses in vivo in mice. *Front Cell Neurosci* **9**, 214, doi:10.3389/fncel.2015.00214 (2015).

8 Abdellah, M. *et al.* Reconstruction and visualization of large-scale volumetric models of neocortical circuits for physically-plausible in silico optical studies. *BMC Bioinformatics* **18**, 402, doi:10.1186/s12859-017-1788-4 (2017).

9 Molineux, M. L., Fernandez, F. R., Mehaffey, W. H. & Turner, R. W. A-type and T-type currents interact to produce a novel spike latency-voltage relationship in cerebellar stellate cells. *J Neurosci* **25**, 10863-10873, doi:10.1523/JNEUROSCI.3436-05.2005 (2005).

10 Roth, A. & Hausser, M. Compartmental models of rat cerebellar Purkinje cells based on simultaneous somatic and dendritic patch-clamp recordings. *J Physiol* **535**, 445-472, doi:10.1111/j.1469-7793.2001.00445.x (2001).

11 Midtgaard, J. Membrane properties and synaptic responses of Golgi cells and stellate cells in the turtle cerebellum in vitro. *J Physiol* **457**, 329-354, doi:10.1113/jphysiol.1992.sp019381 (1992).

12 Solinas, S. *et al.* Computational reconstruction of pacemaking and intrinsic electroresponsiveness in cerebellar Golgi cells. *Front Cell Neurosci* **1**, 2, doi:10.3389/neuro.03.002.2007 (2007).

13 Forrest, M. D. Simulation of alcohol action upon a detailed Purkinje neuron model and a simpler surrogate model that runs >400 times faster. *BMC Neurosci* **16**, 27, doi:10.1186/s12868-015-0162-6 (2015).

14 Llinas, R. & Sugimori, M. Electrophysiological properties of in vitro Purkinje cell somata in mammalian cerebellar slices. *J Physiol* **305**, 171-195, doi:10.1113/jphysiol.1980.sp013357 (1980).

15 Chu, C. P., Bing, Y. H., Liu, H. & Qiu, D. L. Roles of molecular layer interneurons in sensory information processing in mouse cerebellar cortex Crus II in vivo. *PLoS One* **7**, e37031, doi:10.1371/journal.pone.0037031 (2012).

16 Hausser, M. & Roth, A. Dendritic and somatic glutamate receptor channels in rat cerebellar Purkinje cells. *J Physiol* **501 ( Pt 1)**, 77-95, doi:10.1111/j.1469-7793.1997.077bo.x (1997).

17 Kondo, S. & Marty, A. Synaptic currents at individual connections among stellate cells in rat cerebellar slices. *J Physiol* **509 ( Pt 1)**, 221-232, doi:10.1111/j.1469-7793.1998.221bo.x (1998).

18 Chavas, J. & Marty, A. Coexistence of excitatory and inhibitory GABA synapses in the cerebellar interneuron network. *J Neurosci* **23**, 2019-2031 (2003).

19 Houston, C. M., Bright, D. P., Sivilotti, L. G., Beato, M. & Smart, T. G. Intracellular chloride ions regulate the time course of GABA-mediated inhibitory synaptic transmission. *J Neurosci* **29**, 10416-10423, doi:10.1523/JNEUROSCI.1670-09.2009 (2009).

20 Christie, J. M. & Westbrook, G. L. Lateral excitation within the olfactory bulb. *J Neurosci* **26**, 2269-2277, doi:10.1523/JNEUROSCI.4791-05.2006 (2006).

21 Mathy, A. *et al.* Encoding of oscillations by axonal bursts in inferior olive neurons. *Neuron* **62**, 388-399, doi:10.1016/j.neuron.2009.03.023 (2009).

22 Hines, M. L. & Carnevale, N. T. NEURON: a tool for neuroscientists. *Neuroscientist* **7**, 123-135, doi:10.1177/107385840100700207 (2001).

23 Martone, M. E. *et al.* The cell-centered database: a database for multiscale structural and protein localization data from light and electron microscopy. *Neuroinformatics* **1**, 379-395, doi:10.1385/NI:1:4:379 (2003).
